# Molecular basis of FIGNL1 in dissociating RAD51 from DNA and chromatin

**DOI:** 10.1101/2024.07.16.603765

**Authors:** Alexander Carver, Tai-Yuan Yu, Luke A Yates, Travis White, Raymond Wang, Katie Lister, Maria Jasin, Xiaodong Zhang

**Affiliations:** DNA Processing Machines Laboratory, Francis Crick Institute, London, UK; Section of Structural and Synthetic Biology, Department of Infectious Disease, Imperial College London, London, UK; Developmental Biology Program, Memorial Sloan Kettering Cancer Center

## Abstract

Maintaining genome integrity is an essential and challenging process. RAD51 recombinase, the central player of several crucial processes in repairing and protecting genome integrity, forms filaments on DNA. RAD51 filaments are tightly regulated. One of these regulators is FIGNL1, that prevents persistent RAD51 foci post-damage and genotoxic chromatin association in cells. The cryogenic electron microscopy structure of FIGNL1 in complex with RAD51 reveals that the FIGNL1 forms a non-planar hexamer and RAD51 N-terminus is enclosed in the FIGNL1 hexamer pore. Mutations in pore loop or catalytic residues of FIGNL1 render it defective in filament disassembly and are lethal in mouse embryonic stem cells. Our study reveals a unique mechanism for removing RAD51 from DNA and provides the molecular basis for FIGNL1 in maintaining genome stability.

## Introduction

RAD51 recombinase is a key protein involved in homologous recombination, the most faithful repair process for a double-stranded DNA break(*1–3*). RAD51, together with its meiosis specific homologue DMC1, also perform analogous functions in meiotic recombination when parental homologous chromosomes pair and recombine(*4*). The RAD51-ssDNA filament catalyzes the essential processes of homology search, strand invasion and heteroduplex formation(*1, 3, 5*). RAD51 needs to be removed from heteroduplex DNA before homology-directed DNA synthesis and repair/recombination can be carried out. Aberrant loading of RAD51 on ssDNA can lead to multiple strand invasions across chromosomes which in turn can lead to the formation of ultrafine bridges and chromosome instability, a hallmark of cancer(*6, 7*). Furthermore, RAD51 is shown to be essential for DNA replication, especially in dealing with replication stress, and to maintain a stable replication fork(*8, 9*). Significantly, over-expression and accumulation of RAD51 on chromosome has been associated with increased drug resistance in tumor cells and increased genome instability and apoptosis in normal cells(*10*). Consequently, RAD51 activity is tightly regulated. For example, the tumor suppressors BRCA2 and RAD51 paralog complex BCDX2 promote RAD51-ssDNA filament formation and stability while several DNA helicases and translocases such as RAD54, the RecQ family, HELQ, RTEL1, and FBH1 can disassemble RAD51-ssDNA or -dsDNA filaments(*5, 11–16*). FIGNL1 (fidgetin-like 1), an essential gene in mice belonging to the large AAA+ ATPase family, has recently been shown to prevent persistent RAD51 foci formation, prevent replication fork instability and suppress ultra-fine chromosome bridges(*17–20*). Conditional knockout of FIGNL1 in mouse spermatocytes results in massive overloading of RAD51 and DMC1 in these cells(*17, 19*). Recently, FIRRM/FLIP has been shown to form a stable complex with FIGNL1 and together they function in several DNA repair pathways including HR and interstrand DNA cross-link (ICL) repair as well as replication fork protection; and FIGNL1-FIRRM/FLIP can disassemble RAD51 filaments(*19, 21–25*). FIGNL1 has thus been firmly established as an important player in genome maintenance. However, the molecular mechanism of RAD51 modulation by FIGNL1 is currently unknown.

## Results

### FIGNL1 ATPase is essential for RAD51 filament disassembly

Silencing or deleting FIGNL1 has been shown to be embryonic lethal in mice(*17*). Indeed *Fignl1^-/-^* mice die during embryogenesis (**Fig. S1A**) and *Fignl1^-/-^* mouse embryonic stem cells (mESCs) are inviable (**Fig. 1A**, **Fig. S1B-D**). The importance of the enzymatic activity of FIGNL1 has been contradictory. Earlier work indicated that ATPase activity was important for RAD51 foci formation and efficient HR(*18*) but a later study suggested that ATPase activity was not necessary for RAD51 filament disassembly(*26*). In order to understand if and how enzymatic activity of FIGNL1 affects its functionalities, we developed a system to express various FIGNL1 mutants from the *Rosa26* locus in *Fignl1^+/-^* mESCs to ask whether the second endogenous allele could be disrupted (Fig. S1). While *Fignl1^-/-^* cells were viable if they expressed wild-type FIGNL1 from the locus, they were not viable if they expressed the mutant, K456A (K447 in human FIGNL1), in the nucleotide-binding Walker A motif (KA, **Fig. 1A, Fig. S1E**), indicating that nucleotide binding of FIGNL1 is essential.

**Figure 1.**
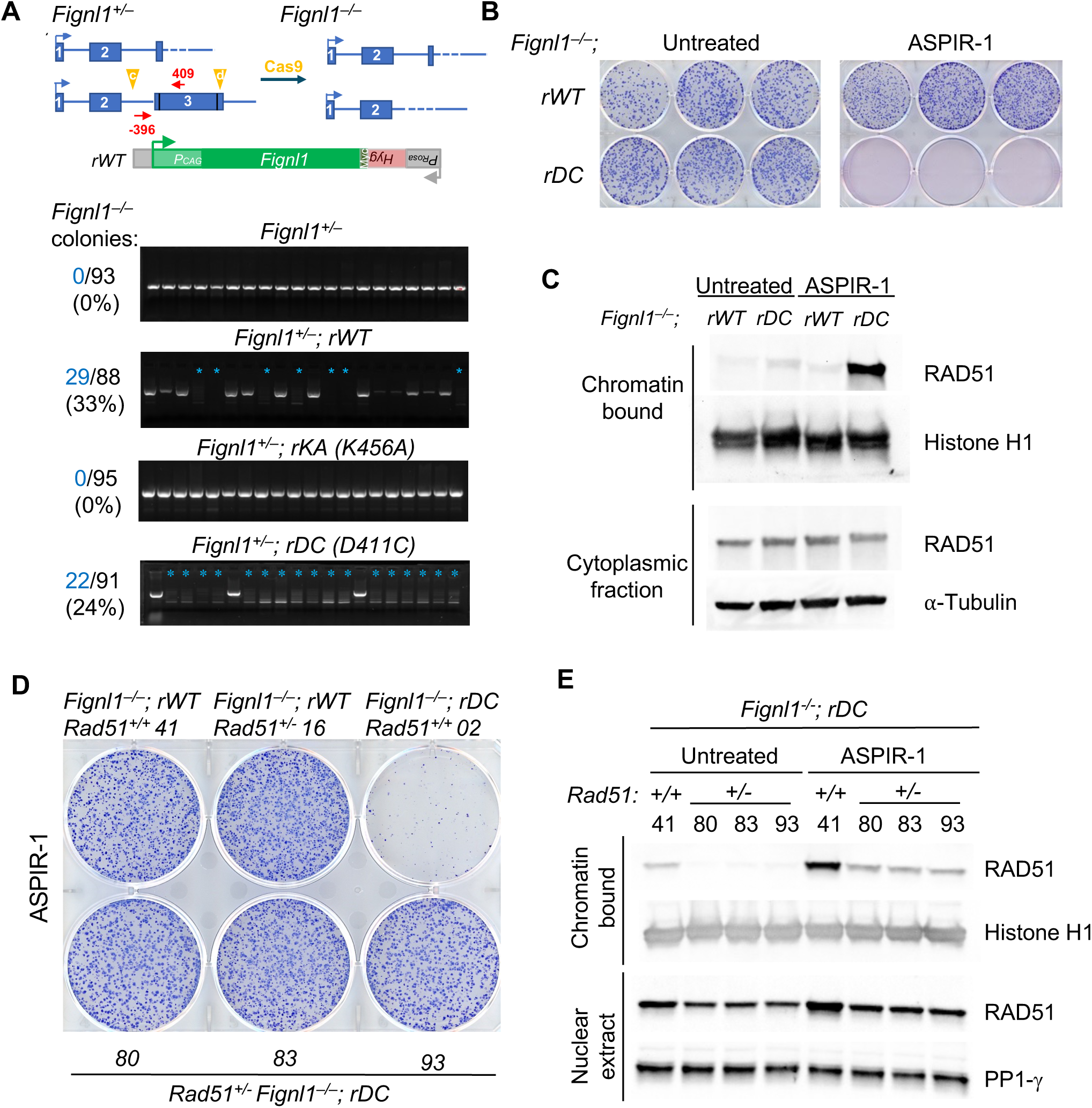
FIGNL1 ATPase activity is required for its functionality in cells. (A) FIGNL1 is required for the viability of mESCs. *Fignl1^+^*^/−^ cells were targeted at the *Rosa26* locus with expression cassettes for wild-type *Fignl1* (*rWT*) or mutants K456A in the ATPase Walker A motif (*rKA*) or D411C for chemical inhibition (*rDC*) and then selected for *Hyg* gene expression from the *Rosa26* promoter. After confirmation of correct targeting, the second *Fignl1* allele was subjected to CRISPR-Cas9 editing using sgRNAs (c+d) to delete the entire *Fignl1* coding region. Genomic DNA was screened by PCR using primers -396 and 409 for the undeleted allele. Successful deletion is indicated by a blue asterisk. The number of *Fignl1*^-/-^ colonies is indicated, along with the total number of colonies that were screened. (B) Chemical inhibition of the FIGNL1 ATPase is lethal. Covalent modification of FIGNL1 at D411C in the ATPase domain by ASPIR-1 impairs colony formation. (C) RAD51 accumulates in the chromatin fraction in *Fignl1*^-/-^; *rDC* but not *Fignl1*^-/-^; *rWT* cells upon ASPIR-1 treatment. Cells are treated with 0.25 µg/ml ASPIR-1 or DMSO for 24 hr and then subjected to chromatin and cytoplasmic fractionation followed by western blot analysis with antibodies to the indicated proteins. (D) *Rad51* heterozygosity rescues survival of the *Fignl1* mutant for colony formation in the presence of ASPIR-1 (0.25 µg/ml) for 7 days. Three independent clones were tested (80, 83, 93). (E) *Rad51* heterozygosity reduces RAD51 chromatin association in the presence of ASPIR-1 to the level found in untreated *Rad51* wild-type cells. Three independent clones were treated with 0.25 µg/ml ASPIR-1 or DMSO for 24 hr and then subjected to chromatin and nuclear fractionation followed by western blot analysis with antibodies to the indicated proteins.

To probe the effects of ATPase activity in cells in a controlled fashion, we used a chemical genetic approach to inhibit the ATPase activity of FIGNL1 by expressing an ATP-analogue sensitive mutant with a cysteine in the active site(*27*) (**Fig. S2A**). This modified version of FIGNL1 (D411C, D402 in human FIGNL1) retains ATPase activity, but when exposed to the compound ASPIR-1, a covalent bond is formed at the active site to abrogate ATPase activity(*27*). We found that while *Fignl1^-/-^* cells ectopically expressing D411C at *Rosa26* were viable, treatment of these cells with ASPIR-1 was toxic (**Fig. 1A-B, Fig. S2B**). Further, ASPIR-1 treatment of D411C-expressing cells resulted in markedly increased association of RAD51 with chromatin (**Fig. 1C, Fig. S2C-D**), which is known to increase chromosome aberrations/aneuploidy(*28*). Strikingly, the lethality caused by *Fignl1* mutation was rescued by *Rad51* heterozygous deletion (**Fig. 1D, Fig. S1F**). Associated with the rescue, cells displayed reduced RAD51 chromatin accumulation (**Fig. 1E**), consistent with recent data showing that RAD51 inhibition could rescue cellular defects due to *Fignl1* knockout(*20*). Together, these data support the notion that FIGNL1 ATPase activity is required for its functionality in the prevention of aberrant RAD51-chromatin association and genome instability.

To understand how FIGNL1’s ATPase activity is linked to modulating RAD51’s association with chromatin, we set out to investigate the molecular mechanisms of FIGNL1-mediated RAD51 dissociation from DNA and filament disassembly *in vitro*. Previous studies have identified three functional domains in FIGNL1: a N-terminal domain containing a largely unstructured region responsible for localisation to DNA damage sites and for interactions with other proteins, followed by the RAD51-binding FRBD (<u>F</u>IGNL1 <u>R</u>AD51-<u>B</u>inding <u>D</u>omain) and AAA+ ATPase domains (**Fig. S3A**)(*18, 19*). The N-terminal domain interacts with FIRRM/FLIP (*21*), and the interactions are important for the mutual stability of FIGNL1 and FIRRM/FLIP(*19, 23–25*). To ensure protein stability in the absence of interacting partners while maintaining its activities for RAD51 modulation, we purified a N-terminal truncated fragment of human FIGNL1 (residues 287-674), equivalent to those used in earlier studies(*18, 26*), that exhibits ATPase activity and RAD51 binding, which we term FIGNL1ΔN (**Fig. S3**). As expected and consistent with previous studies(*17, 22, 26, 29*), FIGNL1ΔN can dismantle RAD51 from both ssDNA and dsDNA (**Fig. 2, Figs. S3-S4**).

**Figure 2.**
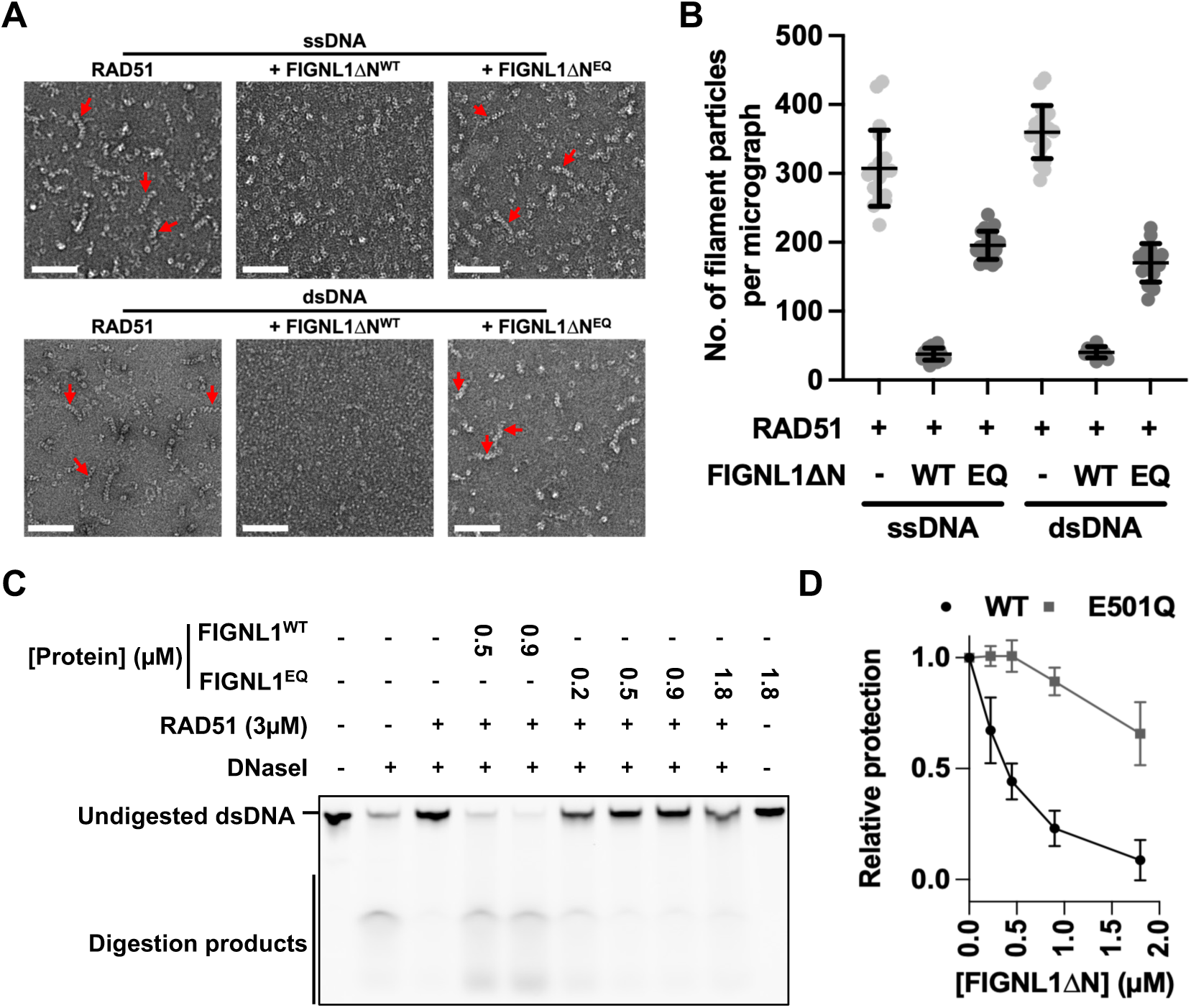
Disassembly of RAD51 filaments is dependent on FIGNL1 ATPase activity. **(A)** Representative negative-stain electron micrographs of RAD51 filaments on both single-stranded and double-stranded DNA in the absence or present of FIGNL1ΔN or FIGNL1DN(E501Q), a mutant in the Walker B motif. Red arrows indicate some of the filaments. Scale bars = 100 nm. **(B)** Quantification of filaments observed in **(A)** indicates a decrease in the number of filaments per micrograph upon incubation with FIGNL1ΔN while FIGNL1DN(E501Q) has little effects. *n* = 18-20 micrographs per condition (**C**) Image of gel, showing nuclease protection of dsDNA by RAD51 in the presence of increasing concentrations of FIGNL1ΔN or E501Q mutant. **(D)** Quantification of nuclease protection assays. n = 3, each data point represents the mean ± s.d. WT FIGNL1ΔN data quantification from independent repeats and 0.5µM and 0.9µM data points shown in **C**. Filament disruption assays shown in **A-D** were carried out using 60nt ssDNA and 60bp dsDNA.

To visualize filament disassembly and to compare the effects of various FIGNL1ΔN mutants in order to probe the requirement of ATPase activities, we imaged and quantified RAD51 filaments using negative stain electron microscopy (NS-EM) (**Fig. 2A-B, Fig. S4**). To assess the activity in solution, we used a nuclease protection assay in which DNA coated by RAD51 is challenged to nuclease treatment(*30*). A decrease in protection indicates dissociation or remodelling of RAD51 from DNA. In the nuclease protection assays, adding FIGNL1ΔN resulted in a decrease in RAD51-mediated protection in a FIGNL1ΔN-concentration dependent fashion **(Fig. 2C-D, Fig. S3C**). Together with the NS-EM results (**Fig. 2A-B**), these data confirm that FIGNL1ΔN disassembles RAD51-DNA filaments. To assess the role of FIGNL1 ATPase activity, we introduced a mutation in the catalytic Walker B motif (E501Q), which is defective in ATP hydrolysis (**Fig. S3B-D**). The E501Q mutant does not affect RAD51 binding (**Fig. S3E**) but is severely defective in disassembling filaments, as judged by NS-EM and nuclease protection assays (**Fig. 2, Fig. S3C, Fig. S4**), suggesting that ATP hydrolysis is required for its activities. Indeed, this mutant resulted in lethality in mESCs (**Fig. S5**). Not surprisingly, the Walker A mutant (K447A) is also defective in ATPase activity and filament disassembly (**Fig. S3B-D**).

Taken together, our cellular and *in vitro* data firmly establish that both nucleotide binding and hydrolysis are required for FIGNL1 to dissociate RAD51 from DNA, filament disassembly and are essential for cell viability.

### Architecture of the FIGNL1-RAD51 complex

Other known ATPases that disassemble RAD51 filaments (collectively termed anti-recombinases) function as DNA translocases or helicases via their ATPase domains(*14, 31*), whereas the AAA+ domain of FIGNL1ΔN does not bind DNA (**Fig. S3F**), suggesting that FIGNL1 acts on RAD51 instead of on DNA. To gain insights into the molecular mechanism of FIGNL1-dependent regulation of RAD51, we utilised FIGNL1ΔN^E501Q^, which forms a stable complex with RAD51 but is defective in filament disassembly, to obtain a structure of FIGNL1 in complex with RAD51, which likely represents an initial engaged state of FIGNL1. Using cryogenic electron microscopy (cryoEM), we determined the structure of the FIGNL1ΔN^E501Q^ -RAD51 complex in the presence of ATP and Mg^2+^ (**Fig. S6**). Single particle analysis produced a 3D reconstruction that revealed a hexameric ring shape, which we attributed to FIGNL1, a AAA+ protein that forms hexamers in the presence of nucleotide (ATPγs) (**Fig. S6E**)(*24*), and additional density above the ring, tethered to the hexameric density, to be RAD51 (**Fig. 3A**). The resolution of this complex is limited, with a global resolution of ∼ 8 Å, likely due to the innate flexibility of the FIGNL1-RAD51 complex (**Fig. S6**). Indeed the identified RAD51-binding FRBD is predicted to be in a largely unstructured region (**Fig. S3A**). Using AlphaFold2(*32, 33*), we generated a model of the FIGNL1 FRBD in complex with a RAD51 trimer, which resembles a filament (**Fig. S7A**). This model predicts that the AAA+ domain is tethered to the FRBD-RAD51 sub-complex via a flexible linker (**Fig. S7A**), consistent with our cryoEM reconstruction (**Fig. 3A**). In this model, the FRBD can bind to a RAD51 trimer using two sites, separated by ∼ 40 amino acids (**Fig. S7A**). The first site (FKTA), which has been previously identified(*18*), is predicted to bind to RAD51 analogous to that of the BRCA2 BRC4 motif (FxxA), that binds at the RAD51 protomer interface in the filament(*34*). A second site (FVPP), which is also present in BRCA2 and RAD51AP1(*35, 36*), binds the adjacent RAD51 protomer via a different location on RAD51 and does not overlap with BRC4 binding site (**Fig. S7A**). Mutating either of the two sites is only moderately defective in our nuclease protection assays while mutating both sites severely reduced its ability in filament disassembly (**Fig. S7B**). Murine ESCs are viable even when both sites are mutated (**Fig. S7C**), suggesting there are other interaction sites between RAD51 and FIGNL1, either directly or via other interacting partners, to partially compensate for the lost interaction site. A recent study has shown that mutating site 1 (FxxA to ExxE) failed to suppress ultra-fine bridge (UFB) formation in U2OS cells(*20*), suggesting a crucial role of this interaction site in UFB suppression.

**Figure 3.**
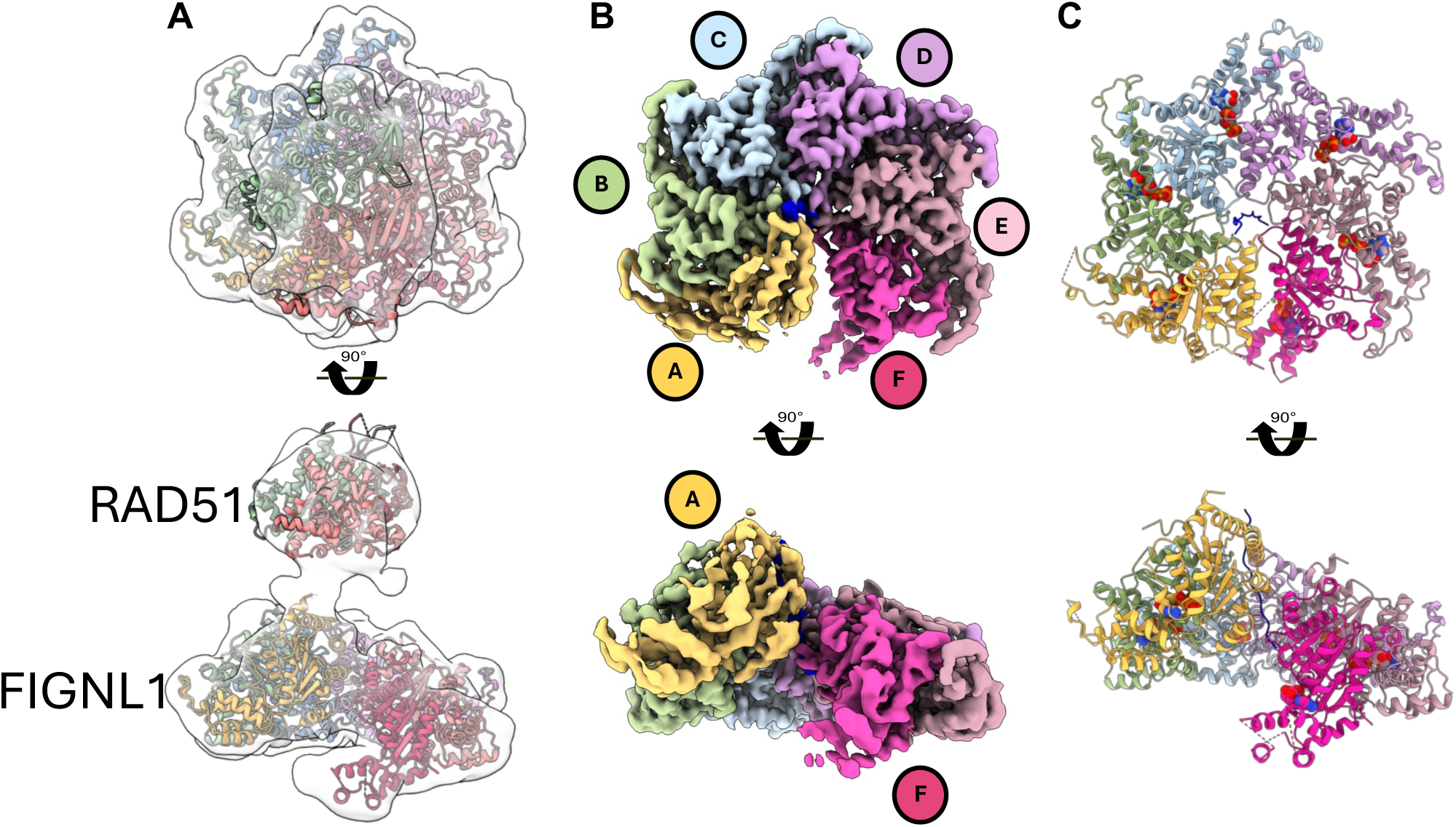
CryoEM structures of the FIGNL1-RAD51 complex and FIGNL1 AAA+ hexamer. **(A)** Top and side views of the cryoEM map and model of the FIGNL1DN^E501Q^-RAD51 complex in the presence of ATP.Mg^2+^. **(B)** Top and side views of the cryoEM map (2.9Å) of the FIGNL1 AAA+ hexamer in the presence of ATP.Mg^2+^. (**C)** Top and side views of the atomic model of the FIGNL1 AAA+ hexamer modelled from the map shown in **B**.

To improve the resolution of the reconstruction, we focused on the FIGNL1ΔN hexamer for further processing, which was resolved to 2.9Å (**Fig. 3B, Fig. S8**). The six FIGNL1 AAA+ domains form a spiral hexamer (**Fig. 3B**), with chain F at the base of the spiral and chain A at the top. The quality of the electron density map is sufficient to allow a structural model of the AAA+ domain of FIGNL1 to be built (**Fig. 3C**, **Table 1, Fig. S8**). We identify clear density for Mg^2+^-ATP, bound in the nucleotide binding pocket of chains A, B, C, D and E, in between adjacent protomers (**Fig. 3C, Fig. S8E**). The electron density of chain F is of poorer quality, presumably due to increased flexibility of this subunit (**Fig. S8B, S8E**). The structural model allowed us to map mutations found in cancer samples and variants of unknown significance in patients with genetic disorders (**Fig. S9A-B**)(*37, 38*). Interestingly, using AlphaMissense (**Fig. S9C, Table 2**) (*39*), E501Q has a high pathogenicity score (0.97) and in human disease mutation database ClinVar, it occurred in 7 samples, consistent with defects we observe *in vitro* and in cells. Several mutations are located at the protomer-protomer interface, and these mutations could interfere with proper assembly of the functional hexamer (**Fig. S9B-C**). Further investigations are required to confirm their potential consequences.

**Table 1.** Cryo-EM data collection, refinement and validation statistics.

|  | FIGNL1-RAD51 complex<br>(EMDB-18946)<br>(PDB 8R64) |
| --- | --- |
| **Data collection and processing** |  |
| Magnification | 81000 X |
| Voltage (kV) | 300 kV |
| Electron exposure (e-/Å<sup>2</sup>) | 50 |
| Defocus range ($\mu\text{m}$) | -0.7 to -2.1 |
| Pixel size ($\text{\AA}$) | 1.1 |
| Symmetry imposed | C1 |
| Initial particle images (no.) | 1.8 million |
| Final particle images (no.) | 57,484 |
| Map resolution ($\text{\AA}$) | 3.2 |
| FSC threshold | 0.143 |
| Map resolution range ($\text{\AA}$) | 2.8 to 5.6 |
| **Refinement** |  |
| Initial model used (PDB code) | 3D8B, Alphafold2 |
| Model resolution ($\text{\AA}$) | 3.4 / 3.1 |
| FSC threshold | 0.5 / 0.143 |
| Model resolution range ($\text{\AA}$) | 2.9 to 3.7 |
| Map sharpening $B$ factor ($\text{\AA}^2$) | -65 |
| Model composition |  |
| Non-hydrogen atoms | 13636 |
| Protein residues | 1774 |
| Ligands | ATP: 5 |
|  | ADP: 1 |
|  | Mg<sup>2+</sup>: 6 |
| $B$ factors ($\text{\AA}^2$) | |
| Protein | 69.2 |
| Ligand | 60.1 |
| R.m.s. deviations |  |
| Bond lengths ($\text{\AA}$) | 0.003 |
| Bond angles ($^\circ$) | 0.53 |
| Validation |  |
| MolProbity score | 1.33 |
| Clashscore | 5.38 |
| Poor rotamers (%) | 0 |
| Ramachandran plot |  |
| Favored (%) | 97.8 |
| Allowed (%) | 2.2 |
| Disallowed (%) | 0 |

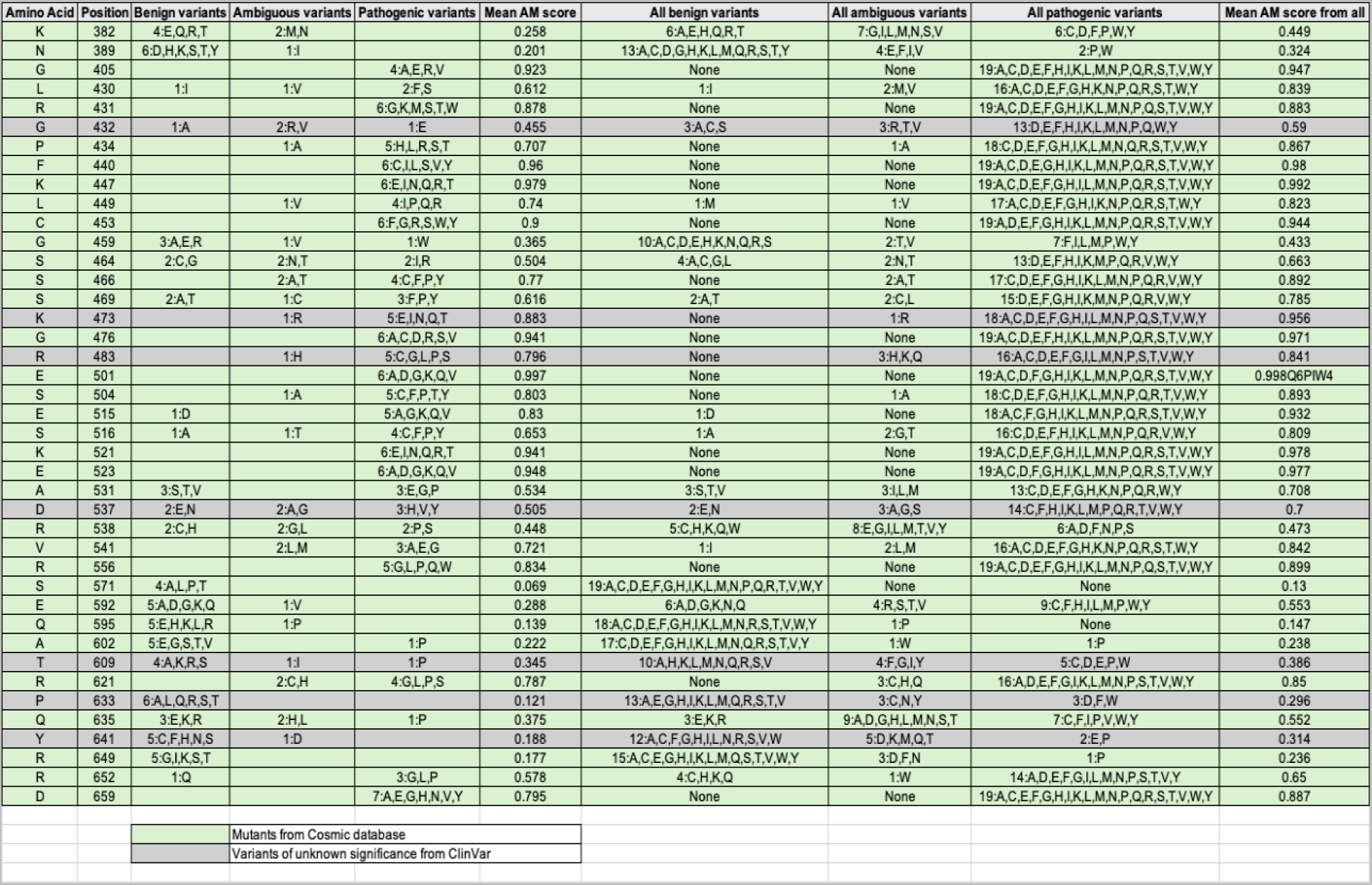

### FIGNL1 coordinates the RAD51 N-terminal peptide in its hexameric pore using pore loops

Unexpectedly, we found additional density consistent with a polypeptide in the central pore of the FIGNL1 AAA+ hexamer (**Fig. 4A**). The polypeptide is tightly coordinated by two highly conserved pore loops of FIGNL1 which form a helical staircase (**Fig. 4B-D**). Many AAA+ proteins utilise pore loops to interact with their respective substrates, either protein polypeptides or nucleic acids(*40–43*). The N-terminal pore loop (residues 470 to 476, pore loop 1, PL1) contains a lysine-tryptophan dipeptide (KW) at its tip which intercalates the side chains of the peptide (**Fig. 4D**). The C-terminal pore loop (residues 506-514, pore loop 2, PL2) is less tightly associated with the peptide, but habours histidine 514 that tracks along the peptide backbone (**Fig. 4C-D**). The extra density in the pore suggests that in addition to the two identified FRBD sites, there are additional interaction sites between RAD51 and FIGNL1. We could fit residues 2-13 of the RAD51 N-terminus, which belong to an unstructured region not observed in previous crystal and cryoEM structures (**Fig. S10A**). The peptide is well resolved, with a local resolution of 2.7 Å (**Fig. S10A-B**). High Q-scores(*44*) not only indicate a good fit of these residues into the density, but the density has a high resolvability, providing further confidence in the residue assignment (**Fig. S10A**). In this structural model, the highly conserved pore loops encircle the physiochemically conserved hydrophobic/polar amphipathic residue pattern found in the N-terminus of RAD51 (**Fig. 4D-E**, **Fig. S10B-E**), via a close network of interactions, which may provide RAD51 sequence preference (**Fig. 4D-E; Fig. S10D-E**).

**Figure 4.**
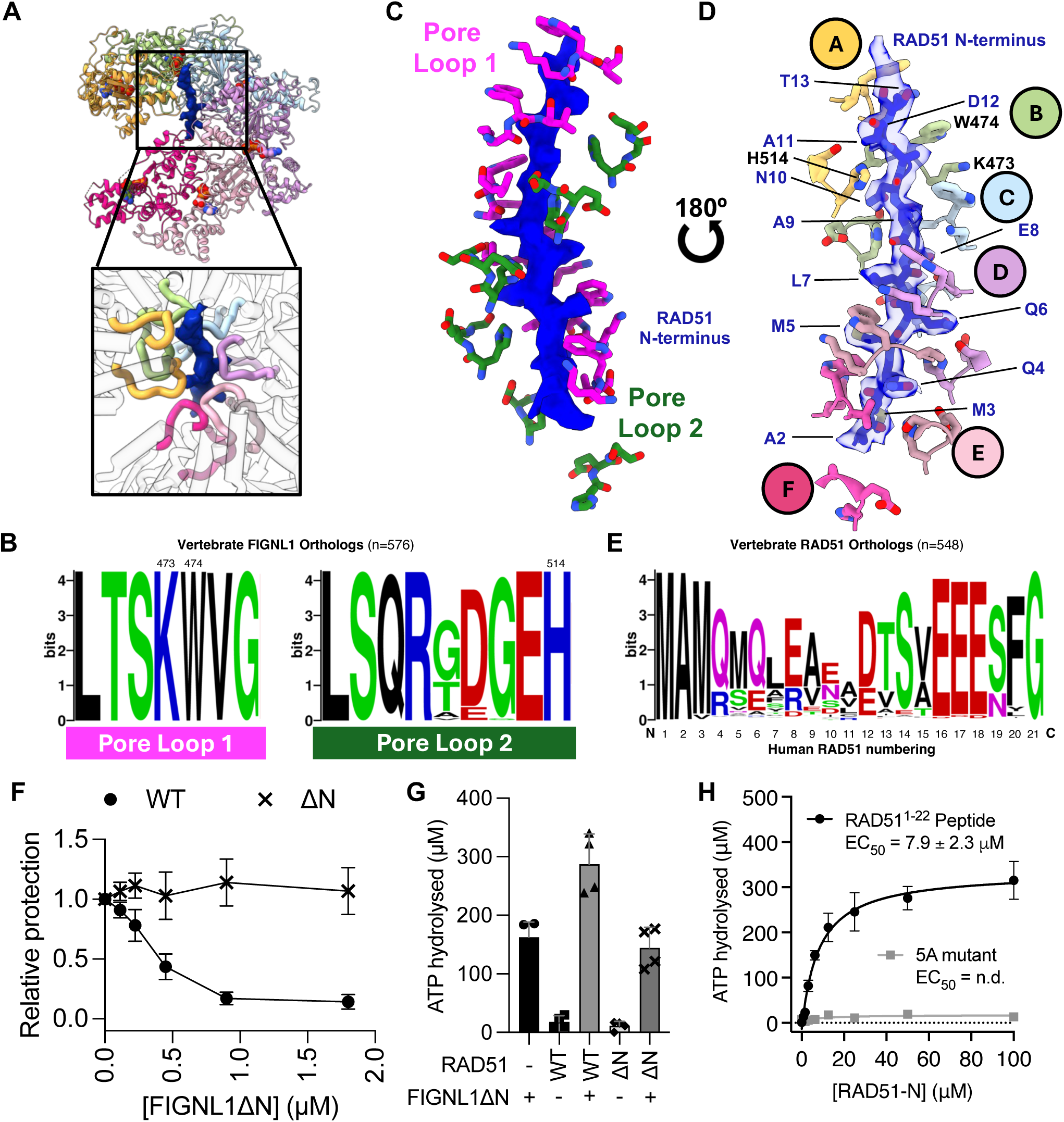
FIGNL1 coordinates the N-terminus of RAD51 through its central hexameric pore. **(A)** Extra density observed in the central pore of the FIGNL1 AAA+ hexamer. **(B)** Sequence conservation plots of the pore loops of FIGNL1 in vertebrates (n=576). Residues 473, 474 and 514 are labelled. **(C)** The pore loops of FIGNL1 form two helical staircases enclosing the RAD51 N-terminus **(D)** The pore loop residues intercalate with the RAD51 N-terminal residues. **(E)** Conservation plot of the N-termini of RAD51 in 548 vertebrates. Numbering refers to the human RAD51 sequence. **(F)** Nuclease protection of ssDNA by RAD51 or RAD51 with N-terminal 20 a.a. deleted (RAD51DN), in the presence of increasing concentrations of FIGNL1DN (**G**) Quantification of ATPase activity of FIGNL1 alone or incubated with RAD51 or RAD51ΔN. (**H**) Dose-response ATPase activity of FIGNL1ΔN in the presence of increasing concentrations of the RAD51 N-terminal peptide or with the first 5 amino acids replaced by alanine (n = 4).

To probe the importance of the RAD51 N-terminus, which has not previously been shown to have specific functions in filament regulation, we investigated if FIGNL1 requires the RAD51 N-terminus for filament disassembly. RAD51 lacking the N-terminal 20 residues (RAD51ΔN) can form filaments and protect DNA from nuclease digestion (**Fig. S11A-C**). Strikingly, RAD51ΔN filaments could not be efficiently disrupted by FIGNL1ΔN in the nuclease protection assay (**Fig. 4F**, **Fig. S11D**). This is not due to the lack of interactions between FIGNL1ΔN and RAD51ΔN, as the binding, although reduced, is still sufficiently high to ensure interactions under the experimental conditions (**Fig. S11E**). This is consistent with the FRBD being the main recruitment site between FIGNL1 and RAD51. Given that ATPase activities of some AAA+ proteins are stimulated by their respective substrates(*41, 45*), we tested whether RAD51 stimulates the ATPase activity of FIGNL1. The ATPase activity of FIGNL1ΔN is enhanced in the presence of RAD51 (**Fig. 4G**) but this enhancement is abolished when the N-terminus of RAD51 is deleted (**Fig. 4G**). Adding FIGNL1ΔN defective in ATPase activity (K447R or E501Q) does not stimulate total ATP hydrolysis, while adding ATPase defective RAD51 (K133R, RAD51-WA) still showed significant enhancement, suggesting the majority of stimulation is due to increased ATPase activity of FIGNL1 in the presence of the N-terminus of RAD51 (**Fig. S12A-B**). A synthetic peptide containing the complete unstructured 22 amino acid RAD51 N-terminus, or the N-terminal 13 or 18 amino acids, can stimulate FIGNL1 ATPase equivalently (**Fig. 4H, Fig. S12C**). This stimulation relies on the very N-terminal amphipathic region of RAD51 observed in our cryoEM structure as replacing the first 5 residues with alanine in this peptide diminishes the stimulation (**Fig. 4H**). Furthermore, FIGNL1 ATPase stimulation is specific to the RAD51 N-terminus, as high concentrations of a peptide of poly-glutamate, tubulin C-terminal tails(*46*), ssDNA, and dsDNA all fail to enhance its ATPase activities (**Fig. S12C-D**). Together these data reveal that FIGNL1ΔN utilises the RAD51 N-terminal region to stimulate its ATPase activity, and that the RAD51 N-terminus is essential for its ability to disassemble RAD51 filaments.

### Pore loop integrity is required for filament disassembly and mESC viability

To corroborate the structural data and to further investigate the importance of the FIGNL1 pore loops, we substituted the KW of pore loop 1 to a glutamic acid and alanine respectively (K473E,W474A, herein referred to as the PL mutant) (**Fig. S13A**). This mutant retains near wild-type ATPase activity and RAD51 binding (**Fig. S13B-C**). This mutant ATPase activity can be stimulated by the RAD51 N-terminus, but to a lesser extent compared to wild-type (**Fig. S13D**). Critically, this mutant is severely defective in disassembling RAD51 filaments on both dsDNA and ssDNA as can be visualised using NS-EM (**Fig. 5A-B**) and nuclease protection assays (**Fig. 5C-D**). The equivalent mutation in mESCs is incompatible with cell viability (**Fig. 5E**). Together these data support a crucial role of the pore loops for FIGNL1 activity.

**Figure 5.**
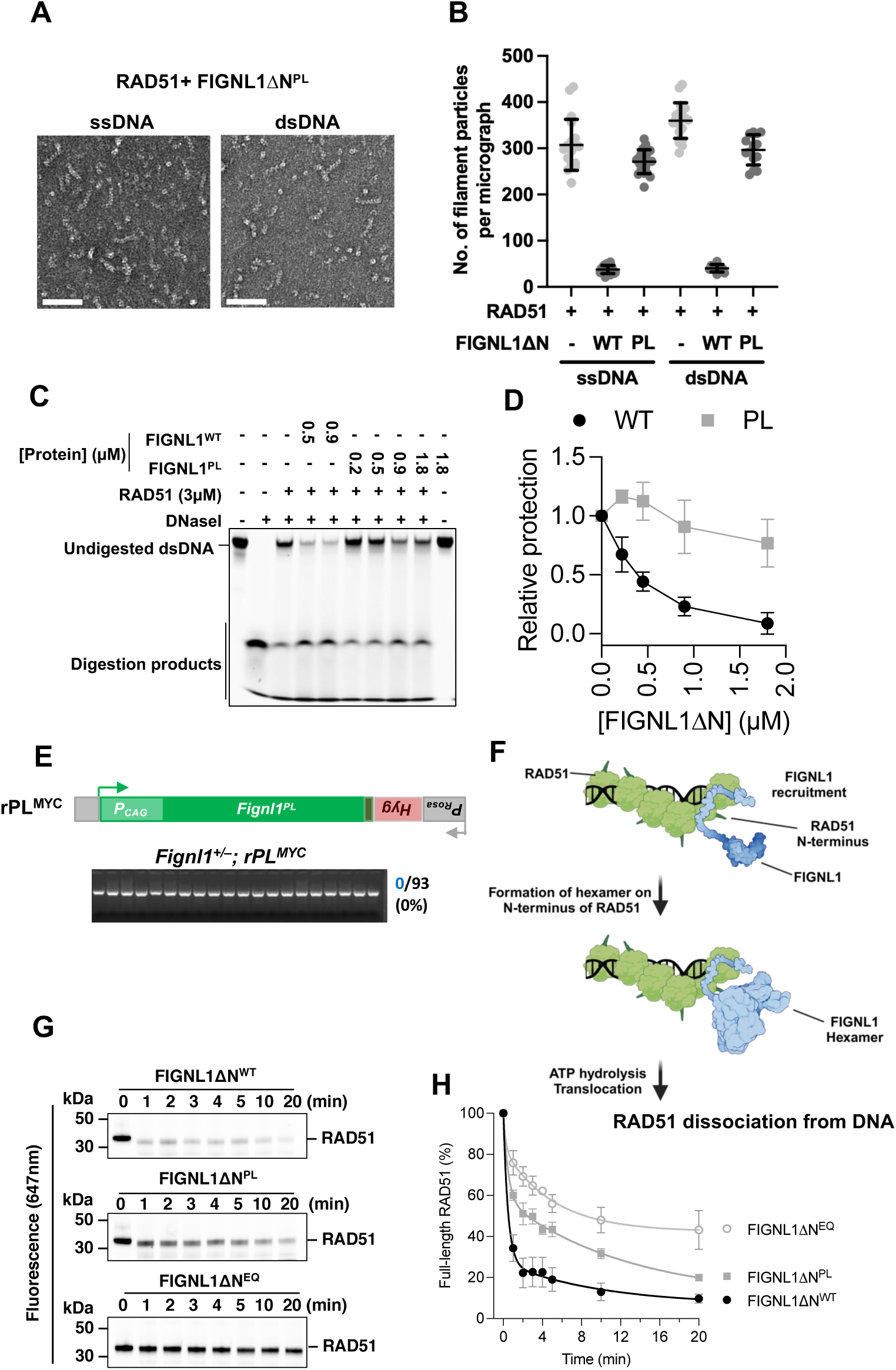
Mutation of pore loop 1 confers loss of RAD51 filament disassembly and cell lethality. **(A)** Representative micrographs of RAD51-DNA filaments treated by FIGNL1ΔN bearing mutations in pore loop 1 (PL mutant) with ss or ds-DNA. Scale bars = 100 nm **(B)** Quantification of experiments shown in (**A)**, highlighting the loss of filament disassembly by FIGNL1ΔN bearing PL mutation. *n* = 20, data are shown as mean ± s.d. WT FIGNL1ΔN data is for comparison and is as shown in Fig. 2. **(C)** Nuclease protection of dsDNA coated by RAD51 upon treatment with wildtype or PL mutant of FIGNL1ΔN. **(D)** Quantification of RAD51 + FIGNL1ΔN^PL^ experiments shown in **c**, *n* = 3, data is shown mean ± s.d. WT FIGNL1ΔN data quantification from independent repeats shown in Fig. 3 and 0.5µM and 0.9µM data points shown in **C**. **(E)** FIGNL1 pore loop mutant K483E/W484A (PL) is not compatible with cell survival. An expression cassette for the mutant (*rPL^Myc^*) was targeted to *Rosa26* locus as in Fig. 1A. No *Fignl1*^-/-^ colonies were obtained after CRISPR-Cas9 gene editing in *Fignl1^+^*^/−^ cells with gRNAs c and d as in Fig. 1A. **(F)** A proposed model of RAD51 filament disassembly by FIGNL1. FIGNL1 is recruited to the RAD51 filament via its FRBD domain and forms a hexamer, enclosing the N-terminus of RAD51, which stimulates the ATPase activity of FIGNL1 and promotes translocation of the N-terminus in the hexamer pore, leading to the unfolding/removing of the RAD51 from the filament, promoting disassembly. (**G**) SDS-PAGE gels showing RAD51 degradation by proteases in the presence of FIGNL1DN and its mutants with ATP. (**H**) quantification of RAD51 band intensity over time.

Recent studies have shown FIGNL1 is important in meiotic DMC1-focus formation and resolution(*17, 19*). We thus set out to test the ability of FIGNL1ΔN in disassembling DMC1 filaments *in vitro*. DMC1 does not efficiently protect dsDNA from nuclease(*47*); we therefore used ssDNA in our assays. FIGNL1ΔN could also reduce nuclease protection by DMC1 (**Fig. S14A-B**), indicating disruption of nucleoprotein complexes, and both DMC1 and DMC1 N-terminal peptide stimulate ATPase activity of FIGNL1ΔN (**Fig. S14C-D**). However the DMC1 N-terminal peptide is ∼3-fold less effective at stimulating FIGNL1 ATPase activity compared to that of RAD51 under the same experimental conditions (**Fig. S14D)**. DMC1 and RAD51 N-terminal regions share some degree of conservation (**Fig. S14E**), suggesting that FIGNL1 substrate selection is somewhat plastic. Importantly, the pore loop mutant is severely defective in disassembling DMC1 filaments (**Fig. S14A**). Together these data suggest that FIGNL1 can also disrupt DMC1-DNA association using a similar mechanism to that of RAD51, despite being less efficient in our assays. The precise mechanism and sequence specificity of FIGNL1 requires further investigation.

## Discussions

### FIGNL1 acts as a peptide translocase to disociate RAD51 from DNA

Our data presented here suggest a molecular mechanism of how FIGNL1 acts on RAD51 during dissociation from DNA and filament disassembly. We showed that FIGNL1 AAA+ domains do not bind DNA and DNA does not stimulate its ATPase activities (**Fig. S3F, S12C**), and thus are unlikely to act as an ATP-dependent DNA translocase. Instead, we show that the ATPase activity is stimulated by RAD51, specifically via the very N-terminus, which we observe is enclosed by the FIGNL1 AAA+ hexamer. Therefore FIGNL1 likely acts directly on RAD51 via interacting with RAD51 N-terminus. The FRBD is responsible for the direct recruitment of FIGNL1 to RAD51. Site 1 (FxxA) binds to RAD51 at a location that overlaps with the protomer interface in a RAD51 filament, suggesting that site 1 would prefer to bind to the 3’-end of a RAD51 filament or a gap in the filament. Site 2 (FVPP), however, could bind to a RAD51 molecule inside the filament, suggesting that FIGNL1 can be recruited to anywhere along the filament. Mutating one of the two sites has mild defects in our nuclease protection assays (**Fig. S7B**). Interestingly, *in vivo*, mESCs with both sites mutated are still viable while those with pore loop mutations are not (**Fig. S6C)**, confirming that FRBD mainly acts in recruitment and that among the three interaction sites between FIGNL1 and RAD51, pore loops enclosing the RAD51 N-terminus are the most important. It is likely that *in vivo*, FIGNL1 could also be recruited to RAD51 via other interacting partners, such as FIRRM/FLIP, which is shown to bind to RAD51(*25*). Our data thus support a model that FIGNL1 is recruited to the filament through its FRBD (and other interacting partners), on the side of the filament. FIGNL1 hexamer assembles around and encloses the RAD51 N-terminus via an intricate network of interactions (**Fig. 5F**). Indeed 3D variability analysis of the FIGNL1-RAD51 structure reveals a mixture of tetrameric, pentameric, and hexameric configurations of the FIGNL1 AAA+ hexamer on the RAD51 N-terminus (**Fig. S15**). This model is also consistent with the observed minimal level of ATP hydrolysis by FIGNL1 at submicromolar concentrations, which is stimulated by RAD51 N-terminus since full ATPase activity requires hexamer formation (**Fig. 4H)**. A similar mechanism has been proposed for Katanin, which oligomerizes as a hexamer on microtubules(*48*).

Thus far all the known AAA+ proteins that enclose substrate peptides in their hexamer central pores utilise ATP hydrolysis to translocate or remodel the substrate(*49*). Given the similarities of the structure of FIGNL1ΔN in complex with RAD51 presented here with that of Vps4, Spastin, Katanin and TRIP13 in complex with their substrate peptides(*40, 41, 50–52*) (**Fig. S16**), we propose that FIGNL1 acts as a peptide translocase that leads to RAD51 remodelling. Using limited proteolysis, we showed that RAD51 was sensitive to Proteinase K digestion in the presence of FIGNL1ΔN and ATP, but markedly less so in the presence of PL or EQ mutants (**Fig. 5G-H**). Proteolytic patterns of RAD51 also differ between FIGNL1 mutants, suggesting that RAD51 conformation is altered by FIGNL1, resulting in the differential proteolytic sensitivities (**Fig. S17**). Indeed, FIGNL1-RAD51 N-terminal interactions strongly resemble that of TRIP13 in complex with the with MAD2 N-terminus (**Fig. S16**). TRIP13 uses its ATPase activity to translocate the unstructured N-terminal region of MAD2 via its pore loops, partially unfolding MAD2 and converting an active closed conformation to an inactivate open conformation (*53, 54*).

Similarly to other AAA+ translocases, FIGNL1 translocates by ATP hydrolysis along the RAD51 N-terminus via the pore loops(*55–57*), leading to pulling or partial unfolding/remodelling of the RAD51 molecule within the filament, destabilising and disassembling the filament as the N-terminal domain links adjacent ATPase domains of RAD51 in the filaments(*58*). Indeed the FIGNL1-PL mutant is defective in filament disassembly, despite stimulated ATPase activities, and does not increase RAD51 sensitivity to proteolysis, supporting the notion that the PL residues are important for its translocating activities analogously to other AAA+ translocases(*41, 52*). Our model suggests that in addition to acting on RAD51 nucleoprotein filaments, FIGNL1 could also act on RAD51 bound to other proteins including nucleosomes, as shown recently(*59*) as the action does not require the acccess of FIGNL1 to DNA.

### FIGNL1’s unique mechanism in dissociating RAD51 from chromatin defines its critical roles in cell viability

The unique mechanism of FIGNL1 in disassembling RAD51 from DNA filaments, acting anywhere along the RAD51 filaments, the potential ability of dissociating RAD51 from other binding partners, including nucleosome-bound RAD51, and the ability to remodel RAD51, suggest FIGNL1 act as a general RAD51 regulator. Our DMC1 data suggests a conserved mechanism in FIGNL1-mediated recombinase filament disassembly. This is distinct from other anti-recombinases which act on DNA from the end of filaments, and may explain the critical role of FIGNL1 in cell viability that cannot be substituted by other anti-recombinases. Excess RAD51 accumulation on ssDNA is predicted to cause abberant HR and interfere with ongoing replication. Furthermore accumulation of RAD51 on dsDNA and chromatin could interfere with other DNA transactions such as chromatin remodelling and transcription. In normal cells, FIGNL1 and other anti-recombinases prevent RAD51 accumulation on chromatin and abberant loading on DNA. Absence of FIGNL1 will result in RAD51 chromatin accumulation, some of which cannot be removed by other recombinases, leading to cell death. Indeed, heterozygous *Rad51* deletion results in reduced RAD51 chromatin association, and rescues viability associated with *Fignl1* mutation. Our work here thus explains the critical roles of FIGNL1 in cell viability and reveals the molecular mechanism of RAD51 filament disassembly and the maintainence of genome stability.

## Acknowledgements

Initial screening of electron microscopy grids was carried out at Imperial College London Centre for Structural Biology. We acknowledge Diamond Light Source for access and support of the cryoEM facilities at the UK national eBIC, proposal EM19865, funded by the Wellcome Trust and MRC, and London Consortium for high resolution cryoEM (LonCEM), funded by the Wellcome Trust. We thank Natalie Jones and Tarun Kapoor for discussions and the gift of ASPIR-1. We thank members of the Zhang and Jasin Labs for their helpful insights and discussions.

## Author contributions

A.C., T.Y., M.J. and X.Z. designed the project. CryoEM sample preparation, image processing and structure determination was performed by A.C. and L.A.Y. Protein construct cloning, mutagenesis, and purification was performed by A.C. with contributions from K.L. and L.A.Y. *In vitro* assays were performed by A.C. and L.A.Y, with contributions from K.L. All cellular work was carried out by T.Y. with assistance from T.W. Mouse studies were carried out by R.W. Bioinformatic analysis was performed by L.A.Y., A.C. and X.Z. wrote the initial manuscript with contributions from L.A.Y., T.Y. and M.J. Edits to the manuscript were carried out by all authors.

## Data Availability

All the data have been deposited in wwPDB with access codes EMDB-18946, PDB code 8R64

## Supplementary Materials

### Materials and Methods

#### Protein expression and purification

A truncation of FIGNL1 (FIGNL1ΔN – amino acids 287-674) was expressed in *Escherichia coli* BL21(DE3) cells as an N-terminal 6xHistidine (HisTag) fusion with a C-terminal single Strep-tag and purified by affinity chromatography (HisTrap followed by StrepTrap) and size exclusion chromatography using a Superdex 200 increase 10/300 GL column (GE Healthcare). FIGNL1ΔN eluted as a single peak which was measured by SEC-MALLS (Wyatt Technology) to be a monomer. The protein was concentrated to ∼1.5mg/mL and stored in FΔN Buffer (20mM Tris pH 8.0, 300mM NaCl, 1mM TCEP, 5% glycerol), flash frozen and stored at -80°C. All subsequent site-directed mutagenesis was performed using the CloneAmp mutagenesis protocol and the proteins were purified as described above. RAD51 was purified as previously described with some alterations(*60*). In brief, RAD51 was co-expressed with 6xHis-MBP-BRC4 (pOPINM) and GroEL (pCh1/RAD51) in *E. coli* BL21(DE3) without tags and purified through affinity chromatography, followed by Heparin affinity chromatography and finally anion exchange. The protein was dialysed against RAD51 storage buffer (20mM Tris pH 8.0, 150mM NaCl, 10% glycerol, 1mM TCEP), aliquoted and stored at a concentration of 1.5 - 2.0mg/mL at -80°C. DMC1 was expressed in *E. coli* BL21 pLysS cells as an N-terminally tagged 6xHistidine fusion protein. A plate of transformants was resuspended in 10mL of LB media and 5mL of this resuspension was added to 2x500mL LB supplemented with Ampicillin and Chloramphenicol (LB-Amp/Cam). The cultures were grown at 37°C to an OD of ∼ 0.5 and then incubated for 1hour at 18°C without shaking. Expression was induced with 0.1mM IPTG and incubated with shaking at 18°C for 18hours. Cells were pelleted at 4,000rpm, resuspended in cold PBS and pelleted again for storage at -80°C. Cells were thawed, resuspended in lysis buffer, sonicated and the lysate clarified at 36,000x g for 1 hour at 4°C. The clarified lysate was filtered and purified by affinity chromatography (HisTrap followed by Heparin affinity) and the tag cleaved by incubation overnight with GST-3C-protease. Peak fractions were further purified by anion exchange (MonoQ). The purest fractions were then pooled, aliquoted, flash frozen and stored at -80°C.

#### Nuclease protection assay

RAD51 filaments were formed on dsDNA by incubating 3μM RAD51 with 75nM 60mer Cy5-labelled dsDNA (OLIGO1+OLIGO2) in the presence of NP buffer (35mM Tris pH 7.5, 25mM KCl, 2.5mM MgCl2, 1mM TCEP, 1mM ATP) and incubated for 5 minutes at 37°C. FIGNL1ΔN (and mutants) were added to a final concentration of 225nM, 450nM, 900nM and 1.8μM and incubated for 10 minutes at 37°C. 2 units of DnaseI (NEB) was then added to samples and incubated for a further 10 minutes at 37°C. 2μL of NP buffer (without ATP) was added to non-DnaseI-treated samples. Samples were deproteinised with Proteinase K in stop solution (40mM Tris pH 7.5, 100mM EDTA, 6% SDS) for 10 minutes at 37°C. Digestion products were separated on a 10% polyacrylamide 1x TBE gel at 110V for 60 minutes at room temperature. Cy5 signal was imaged using a BioRad ChemiDoc imaging system. Gels were analysed in Fiji (ImageJ) and relative protection was calculated in Excel (Microsoft), plotted and analysed in Prism (GraphPad).

#### RAD51 DNA binding electrophoretic mobility shift assays (EMSAs)

Binding of RAD51 and RAD51ΔN to HR-related DNA substrates was performed at room temperature. DNA substrates(*61*) were ordered from IDT (Table 2) and annealed in annealing buffer (Tris, MgCl2). 15nM of DNA substrate was incubated with increasing concentrations of RAD51 or RAD51ΔN (0.2-5μM) in EMSA buffer (20mM Tris, 100mM NaCl, 1mM TCEP, 1mM ATP, 5mM MgCl2) for 10 minutes at 25°C. Protein-DNA species were separated by gel electrophoresis using 3-12% Bis-Tris NATIVE-PAGE gels (ThermoFisher Scientific) at room temperature for 70 minutes at 60V in 1x TAE running buffer. Protein-DNA complexes were visualized using BioRad ChemiDoc and Cy5 settings.

#### Microscale thermophoresis (MST)

RAD51 was labelled with cysteine reactive RED-MALEIMIDE using the manufacturers’ instructions (NanoTemper Technologies). Labelling was confirmed by SDS-PAGE and spectrophotometric analysis. 100nM RAD51 was incubated with a serial dilution of FIGNL1ΔN (and mutants) in MST buffer (20mM Tris pH 8.0, 100mM NaCl, 1mM ATP, 5mM MgCl2, 0.05% TWEEN-20) for 20 minutes at 25°C. MST experiments were performed using a Monolith NT.115 instrument (NanoTemper Technologies) using premium-treated capillaries at 25°C. Binding data were analysed and plotted using GraphPad Prism, with dissociation constants (Kd) calculated in GraphPad Prism. For FIGNL1-RAD51/RAD51ΔN samples, purified FIGNL1ΔN with an N-terminal YBBR-tag (DSLEFIASKLA) was labelled as previously described(*62, 63*). 100nM labelled FIGNL1ΔN was used for MST experiments and was incubated with a serial dilution of RAD51 and RAD51ΔN and measurements taken as in the labelled RAD51 experiments.

#### ATPase activity assays

ATPase activities of FIGNL1ΔN (and mutants) and RAD51 were measured using the Malachite Green Phosphate Assay Kit (Sigma-Aldrich) according to manufacturers’ instructions. For base ATP hydrolysis of WT FIGNL1ΔN vs PL and E501Q mutants, 3µM of FIGNL1ΔN was incubated with 1mM ATP in ATPase buffer (30mM Tris-HCl pH7.5, 100mM KCl, 20mM NaCl, 3.5mM MgCl2, 5% glycerol) for 30 minutes at 37°C. The samples were diluted 1:20 in ultrapure water and 40µL of reaction was added to 10µL working reagent. For FIGNL1ΔN stimulation by RAD51 and RAD51ΔN, 3μM FIGNL1ΔN or 3µM RAD51 was used, and the reaction was carried out as before. For the peptide stimulation assay, 100nM FIGNL1ΔN was incubated with indicated concentrations of peptide in reaction buffer. Samples were diluted and A620 measured as before. Samples were incubated in the dark for 30 minutes at 25°C and absorbance at 620nm measured in a Clariostar Plate Reader (BMG Labtech). Assays were repeated in triplicate. Free phosphate was calculated by plotting a standard curve of the A620 of known phosphate concentrations. Data was plotted and analysed using Prism (GraphPad).

#### Electron microscopy filament disassembly assay

RAD51 (1µM) filaments were formed on 50nM 60mer ssDNA and dsDNA in the presence of 1mM ATP and 5mM MgCl2 in TBS100 for 5 minutes at 37°C. Filaments were challenged with TBS100 (filament alone) or 300nM WT/E501Q/PL FIGNL1ΔN and incubated for a further 10 minutes at 37°C. 4µL of sample was applied to a 300mesh carbon coated copper grid (AgarScientific) glow-discharged in air for 30 seconds on high (Harrick Plasma Cleaner). After blotting and washing of the grid with two 20µL drops of MilliQ water, the grid was stained with 2% uranyl acetate and blotted to remove excess. Grids were imaged at 29k magnification (pixel size 3.1Å) on a Tecnai T12 TEM equipped with a Gatan Rio16 camera. 20 micrographs were collected for each condition. Particles were picked using Gautomatch (insert reference) using a particle diameter of 180Å and interparticle distance of 60Å. Particles were extracted in a 100-pixel box and subjected to two rounds of 2D classification. The best filament like 2D class averages were selected and particles manually inspected on each micrograph. Number of filament-like particles were plotted on GraphPad Prism and subjected to One-Way Anova with multiple comparisons analysis.

#### Sample preparation for cryogenic electron microscopy (cryoEM)

10μM of purified FIGNL1ΔN(E501Q) was incubated with 10μM purified RAD51 and incubated in the presence of 1mM ATP and 5mM MgCl2 in EM buffer for 10 minutes at 37°C. The sample was then spun at 16,000g for 15 minutes at 4°C and 4μL was loaded onto a glow discharged (10s, high power, Harrick Plasma Cleaner) Quantifoil R1.2/1.3 300 mesh copper grids. After 30 second wait time and 2 second blot, the sample was plunged into liquid ethane by a Vitrobot Mark IV (Thermo Fisher Scientific). Grids were screened on a GLACIOS Cryo-TEM (Thermo Fisher Scientific) for ice quality and particle distribution. For the FIGNL1ΔN-ATPγs sample, 5µM FIGNL1ΔN was incubated with 1mM ATPγs and 5mM MgCl2 for 15 minutes at 37°C in EM buffer. 4µL of the sample was applied to a glow discharged (same method as for FIGNL1ΔN-RAD51 sample) lacey carbon with an ultrathin layer of carbon copper grid.

#### CryoEM data acquisition

All cryo-EM data were acquired on a Titan Krios IV TEM equipped with a Gatan K3 camera using EPU software (Thermo Fisher Scientific). All image pre-processing (motion correction and CTF estimation) was performed using the RELION automatic processing pipeline (**Table1**). 15,406 movies were collected at a magnification of 80,000 x at a physical pixel size of 1.1Å/pixel with a dose of 50 e-/Å^2^ and a nominal defocus range from -0.7 to -2.1 μm. 500 movies were collected using the same parameters as the FIGNL1ΔN-RAD51 sample for the FIGNL1ΔN-ATPγs sample.

#### Image processing of FIGNL1ΔN-ATPγs planar hexamer

500 movies were motion corrected and CTF corrected in cryoSPARC(*64*). Particles were picked using blob picker and extracted in a box size of 160 pixels down-sampled to 80 pixels (2.2Å per pixel). Particles were subjected to extensive 2D classification followed by ab-initio model generation. The particles within the best model (24,000 particles) were transferred to RELION 4.0(*65*) and subjected to 3D refinement followed by post-processing with a B-factor of -50. The final map had a resolution of 9.2Å.

#### Image processing of FIGNL1ΔN-RAD51 complex

Motion corrected movies (using MOTIONCOR2(*66*)) were also CTF corrected (using Patch CTF estimation) in cryoSPARC as well as in RELION (using CTFFIND-4.1(*^67^*)). Initial processing was carried out in cryoSPARC(*64*). Templates were generated from a 3D map obtained from data collected on the GLACIOS microscope during sample screening. These templates were used for particle picking and particles were extracted with a box size of 300 pixels down-sampled to 100 pixels. After 2D classification in which classes were chosen according to the resolution of the density beneath the FIGNL1 hexamer, four initial 3D models were generated and subjected to heterogenous refinement. The best class was then subjected to homogenous refinement to produce a map of 7Å. These particles were then transferred to RELION and re-extracted in the same box size down-sampled to 100 pixels. An initial model was generated, and a consensus refinement map produced. 3D classification without alignment of just the extra density was performed. The particles with the best resolved extra density were chosen, an initial model was generated and refined to produce a map of ∼9 Å. After post-processing, this map improved to 7.7Å(*65*).

#### Image processing of FIGNL1ΔN-RAD51 complex focussed on hexamer

The same micrographs were used for FIGNL1ΔN-RAD51 and FIGNL1ΔN-peptide processing pipelines. Particles were initially picked using Topaz(*68*) and extracted in a box size of 192 pixels. These particles were extracted 3 times binned (3.3Å/pixel) in a box size of 192 pixels down-sampled to 64 pixels. The RELION pipeline was run on half the dataset from which a clear hexameric map was produced from initial model generation. After 3D classification without alignment (T=4), a clear hexameric volume was produced and refined to ∼ 7Å. These particles were re-extracted in a 192 pixel box down-sampled to 96 pixels and refined to produce a map of ∼5Å. This map was imported into cryoSPARC, and templates were generated. These templates were used for automatic picking using the cryoSPARC template picking tool. Using a low threshold for picking (1000 maximum local minima considered), an average of 725 and 632 particles were picked per micrograph for top and side views respectively. These particles were extracted in a box size of 192 pixels down-sampled to 64 pixels. Extensive 2D classification to remove noise and particles on carbon resulted in 470,000 good particles which were used for a consensus refinement using the 4.2Å map generated in RELION. 3D classification produced 4 classes (275,000 particles) with clear high-resolution features, and these particles were re-extracted, unbinned and transferred to RELION where they were re-extracted in a box size of 192 pixels. Bayesian polishing within RELION was used to measure per-particle motion of this selection of particles(*69*). After refinement and post-processing, a 3.2Å map was achieved; however, due to flexibility within chains A and F, these were less well resolved. After another round of 3D classification with high tau factor (T=20), the best hexameric class was refined and post-processed with a B-factor of -25 to give the final map at a global resolution of 3.0Å. This map provided the most featured RAD51 N-terminus density. Polished particles were also processed in cryoSPARC v4.0 and subjected to homogeneous refinement followed by non-uniform refinement. This yielded a 2.9 Å map of the FIGNL1 hexamer bound to the RAD51 N-terminus. 3D variability analysis was carried out in cryoSPARC v4.0.

#### Atomic model building and refinement

A model of the FIGNL1 AAA domain hexamer was generated using AlphaFold2^21^. This model was then split into the component protomers and fit into the generated map and subjected to real space refinement in Phenix. The model was visually inspected, and manually corrected in Coot. The peptide, ATP and magnesium ions were built *de novo* in Coot. The model was subject to real space refinement in Phenix using secondary structure restraints, ligand restraints, and NCS, in early iterations. This process was performed for several iterations and the final model, without NCS, was checked using MolProbity (**Table1**).

### Bioinformatic analysis

Vertebrate FIGNL1 or RAD51 ortholog protein sequences were retrieved from the NCBI (National Center for Biotechnology Information) database and aligned using COBALT (constraint-based multiple sequence alignment tool). The conservation plots were produced using the MSAs in WEBLOGO(*70*).

### RAD51 limited proteolysis

Limited proteolysis was performed by incubating RAD51, labelled with cysteine reactive RED-MALEIMIDE (NanoTemper Technologies), with 1mM ATP in 30mM Tris-HCl pH7.5, 100mM KCl, 20mM NaCl, 3.5mM MgCl2, 5% glycerol for 10 minutes at 37°C. FIGNL1ΔN (or EQ and PL mutants) was added at a 6:1 M ratio, and the reaction incubated for a further 10 minutes at 37°C. The final concentrations of proteins in the reaction was 3µM of FIGNL1ΔN and 500nM RAD51. Proteinase K (NEB) was added to a final concentration of 1.5nM (1:2000 ratio of ProK:FIGNL1ΔN) and incubated at 25°C to limit the rate of proteolysis. Aliquots were withdrawn at time intervals and stopped by addition of 2x concentrated SDS-PAGE loading buffer and heated to 95°C. The digestion products were separated by SDS-PAGE and RAD51 bands visualised using in-gel fluorescence 647nm. Replicated experiments were quantified using Image Lab software (Bio-Rad) and fraction of full-length RAD51 remaining was plotted over time.

#### Generation of *Rosa26* knock-in *Fignl1* mutant cell lines

A multi-step approach was used to knock out *Fignl1* in mESCs (129/B6 *Blm^tet/tet^)*(*71*). The first step was to create *Fignl1*^+/-^ cells using the dual Cas9/sgRNA expression vector pSpCas9(BB)-2A-Puro (PX459, Addgene 48139) containing gRNA-a 5’-CCATGTCTGTGGAGACGACT-3’ or gRNA-b 5’-GCTGTGTGGAAACTCATGCA-3’.

Both gRNAs were transfected to obtain *Fignl1*^+/-^ colonies. Genotyping was performed using the following PCR primers: mFignl1-396F (-396) 5’-GTGCTAGGGATCAAACACTAGGGT-3’ and mFignl1+2907R 5’-AGCAAAGGAGCTGGCTAACTGTGC-3’ under the following conditions: 95° C, 3 min; 35 cycles of 95° C, 30 s; 60° C, 30 s and 72° C, 1 min; final extension at 72° C, 5 min. PCR products were run on a 2% agarose gel to identify knockout alleles (∼500 bp). Clone het-02 (*Fignl1*^+/Δ2833^) was used in the next step to generate *Rosa26* knock-in *Fignl1* mutants. pBS-Rosa26 targeting vectors for wild-type, K456A ATPase mutant, K483E W484A pore loop mutant, or D411C for chemical inhibition, *Fignl1* cDNA were constructed and flanked by loxP sites with a C-terminal Myc tag. The targeting vectors consist of expression cassette under the control of the pCAGGS promoter and a promoterless *Hyg* gene which is driven by the endogenous *Rosa26* promoter upon correct integration. The *Fignl1* and *Hyg* are flanked by ∼750 bp homology arms to guide targeting to the *Rosa26* locus following induction of a Cas9-mediated DSB, gRNA sequence 5’-TGCTGTCTGAGCAGCAAC-3’ and cells selected with 150 μg/ml hygromycin(*72*). Multiple colonies were picked and confirmed by several genotyping steps(*72*). The final step was to use the *Fignl1*^+/-^*Rosa26* targeted cells to generate cells that were knocked out at the second *Fignl1* endogenous allele, transfecting PX459 plasmids containing gRNA-c 5’-GAGGAGGGAAATAGGGCAAT-3’ and gRNA-d 5’-CCTGCCTCAAAAGATAAGTG-3’.

Genotyping for *Fignl1*^-/-^ was done using the following two sets of PCR primers: mFignl1-3108F 5’-GACCTTGCTTCACTTGGAACTCGGCC-3’ and mFignl1+2907R 5’-AGCAAAGGAGCTGGCTAACTGTGC-3’ to detect the deletion allele product (∼700 bp) or mFignl1-396F 5’-GTGCTAGGGATCAAACACTAGGGT-3’ and mFignl1+409R 5’-CTACTGATGCATCAGCAGGTTCCA-3’ to identify products from alleles that were wild type or had not undergone the full deletion between the two gRNAs (∼805 bp).

To generate *Rad51*^+/-^ cell lines, two gRNAs were expressed along with Cas9 to delete *Rad51* exon 5, which encodes the Walker A motif, disrupting the *Rad51* reading frame in *Fignl1^-/-^; rWT* or *Fignl1^-/-^; rDC* cells: Cas9: Addgene #41815; sgRNAs: Addgene #41824 with gRNA-C87: 5’-ACCTGTAAACCAGCTATGT-3’ and gRNA-C90: 5’-GGCTACTCTTCCACAGGACT-3’.

Genotyping for *Rad51* heterozygosity was performed using the following PCR primers: mRad51-C: 5’-TTATCTAAGTAGGATTCTACCATTGAGC-3’ and mRad51-D: 5’-AGACATAAAGTGTAGACTGTGCAAC-3’ to detect the deletion allele product (∼330 bp) or wild-type (∼537 bp).

To generate the dual *Rosa26* knock-in cell lines, a hygromycin sensitive *Fignl1^-/-^; rDC* clone already expressing a Myc-tagged FIGNL1-DC protein from one *Rosa26* allele was generated by tranfecting the Cas9 expression vector with gRNA expression vector for gRNA-Hygro-2: 5’-GTCCGACCTGATGCAGCTCT -3’. Clone “F” was then used in the next step to generate dual *Rosa26* knock-in *Fignl1* mutants, using pBS-Rosa26 targeting vectors to express FIGNL1 wild-type, K456A Walker A mutant, or E510Q Walker B mutant with an N-terminal FLAG tag. CRISPR-Cas9 targeting to the second *Rosa26* locus used gRNA 5’-TGCTGTCTGAGCAGCAAC-3’. Cells selected with 150 μg/ml hygromycin. Multiple colonies were picked and confirmed by several genotyping steps, including loss of the *Rosa26* wild-type PCR product with primers: Rosa26-5’-out-F: 5’-CCATCACAGTTTGCCAGTGATAC-3’ and Rosa26-3’-out-R: 5’-TAAAAAGTCATTCCACAGTTTGAC-3’ (*72*). The FLAG-tagged specific allele is checked by PCR primers mFL1-VE-F: 5’-GTGGAAAGTGAGGTACCGGTTCGAG-3’ and hyg: 5’-CTGTGTAGAAGTACTCGCCGATAG-3’.

#### Clonogenic survival assay

All mouse ES cell lines were cultured in DMEM-HG medium supplemented with 12.5% stem cell grade FBS (Gemini), 1X Pen-Strep, 1X MEM/NEAA, 1X L-Gln, 833 U/ml LIF (Gemini Cat # 400-495), and 0.1 mM β-mercaptoethanol. Culture dishes were pre-coated with 0.1% gelatin. For clonogenic survival assays, 3000 cells were plated in a well of 6-well plate for one day attachment. Where indicated, 1.5 μg/ml or 3 μg/ml ASPIR-1 (a gift of Tarun Kapoor, Rockefeller University)(*27*) was added to cells, and they were allowed to grow for a week. Alternatively, 0.25 μg/ml ASPIR-1 was added for either one day or two days; after washed out of any residual drug, fresh media was added back for a week recovery. Plates were then fixed in methanol and stained with Giemsa.

#### Cell fractionation and western blotting

For cell fractionation, 4 million cells were plated in a 10-cm plate with 1 μg/ml ASPIR-1 or untreated media for 24 hr. Ten million adherent cells were collected and then washed with ice-cold PBS. Subcellular protein fractionation followed with a kit (Thermo #78840). Briefly, ice-cold 200 μl cytoplasmic extraction buffer containing protease inhibitors was added to the cell pellet, the tube was incubated at 4°C for 10 minutes with gentle mixing and then centrifuged at 500 × g for 5 min. Ice-cold membrane extraction buffer (200 μl) containing protease inhibitors was added to the pellet and then the tube was vortexed for 5 sec, incubated at 4°C for 10 min with gentle mixing, and centrifuged at 3000 × g for 5 min. Ice-cold nuclear extraction buffer containing protease inhibitors (100 μl) was added to the pellet, and then the tube was vortexed for 15 sec, incubated tube at 4°C for 30 min with gentle mixing, and then centrifuged at 5000 × g for 5 min. The supernatant is the nuclear extract fraction. The chromatin-bound fraction was obtained by adding 100 mM CaCl2 and 300 units micrococcal nuclease with 100 μl room temperature nuclear extraction buffer with protease inhibitors, vortexed for 15 sec, incubated at 37°C water bath for 5 min, vortexed for another 15 sec, and then centrifuged at 16000 × g for 5 min to obtain supernatant. To perform western blotting, the two supernatants with 1x SDS loading dye was loaded onto a precast SDS-PAGE gel (Bio-Rad #4561033), transferred onto nitrocellulose membrane, and blocked with 5% milk in PBST (50 mM Tris [pH 7.5], 150 mM NaCl, 0.05% Tween 20) for 1 h. For immunodetection, the following antibodies were used: anti-histone H1 (Cat # sc-8030) from Santa Cruz; anti-Rad51 (Cat # PC-130) from Millipore; anti-Myc (Cat # 2276) from Cell Signaling; anti-PP1γ (Cat # SC-515943 from Santa Cruz).

#### CRISPR-CAS9 knockout mice

Cas9 RNPs with gRNA-a: 5’-CCATGTCTGTGGAGACGACT-3’ and gRNA-d: 5’-CCTGCCTCAAAAGATAAGTG-3’were injected into one cell embryos, which were then implanted into female mice. Two Δ2540 founders which deleted the *Fign1* coding region starting from codon 11 were obtained. *Fignl1*^+/Δ2540^ X *Fignl1*^+/Δ2540^ crosses were set up to attempt to generate *Fignl1*^Δ2540/Δ2540^ knockouts, although none were obtained at weaning or at embryonic day 12.5. Genotyping was performed using the following PCR primers for the Δ2540 allele: mFignl1-24F (-24) 5’-GTCTTGTGTTATAGAACCTTTGACATGGAG -3’ and mFignl1+2907R (2907) 5’-AGCAAAGGAGCTGGCTAACTGTGC-3’ to obtain a 391 bp product and for the wild-type allele: mFignl1-396F (-396) 5’-GTGCTAGGGATCAAACACTAGGGT-3’ and mFignl1+197R (197) 5’-GAACAGGTTGGTAGCAAAGACCTG-3’ to obtain a 593 bp product. PCR was performed separately for the two alleles under the following conditions: 95° C, 3 min; 35 cycles of 95° C, 30 s; 60° C, 30 s and 72° C, 1 min; final extension at 72° C, 5 min.

#### Conditional knockout system

A *Cre*-expression plasmid (Addgene #125821) was transfected into *Fignl1^-/-^rWT (*clone #20). To select for cells with *Cre* expression, 1 µg/ml puromycin added to the cells for one day. After selection, cells were reseeded from 10 cm plates to 6 wells and genomic DNA was collected every 3 to 4 days at passage. Genotyping for *rWT* was done using the following PCR primers: hm-F 5’-CGGTTCGAGGAATTCAGATCTTTTTCCCTCTGCC-3’ and hyg 5’-CTGTGTAGAAGTACTCGCCGATAG-3’ to identify products from *Rosa26* alleles that were wild type or had not undergone the full deletion (∼773 bp). Genotyping for *rΔ* was done using the following PCR primers: Pcag-F 5’-GCAACGTGCTGGTTATTGTGCTGT-3’ and hyg to identify products from *Rosa26* allele that was undergone *Cre-*mediated deletion (∼393 bp).

---

**Fig. S1.**
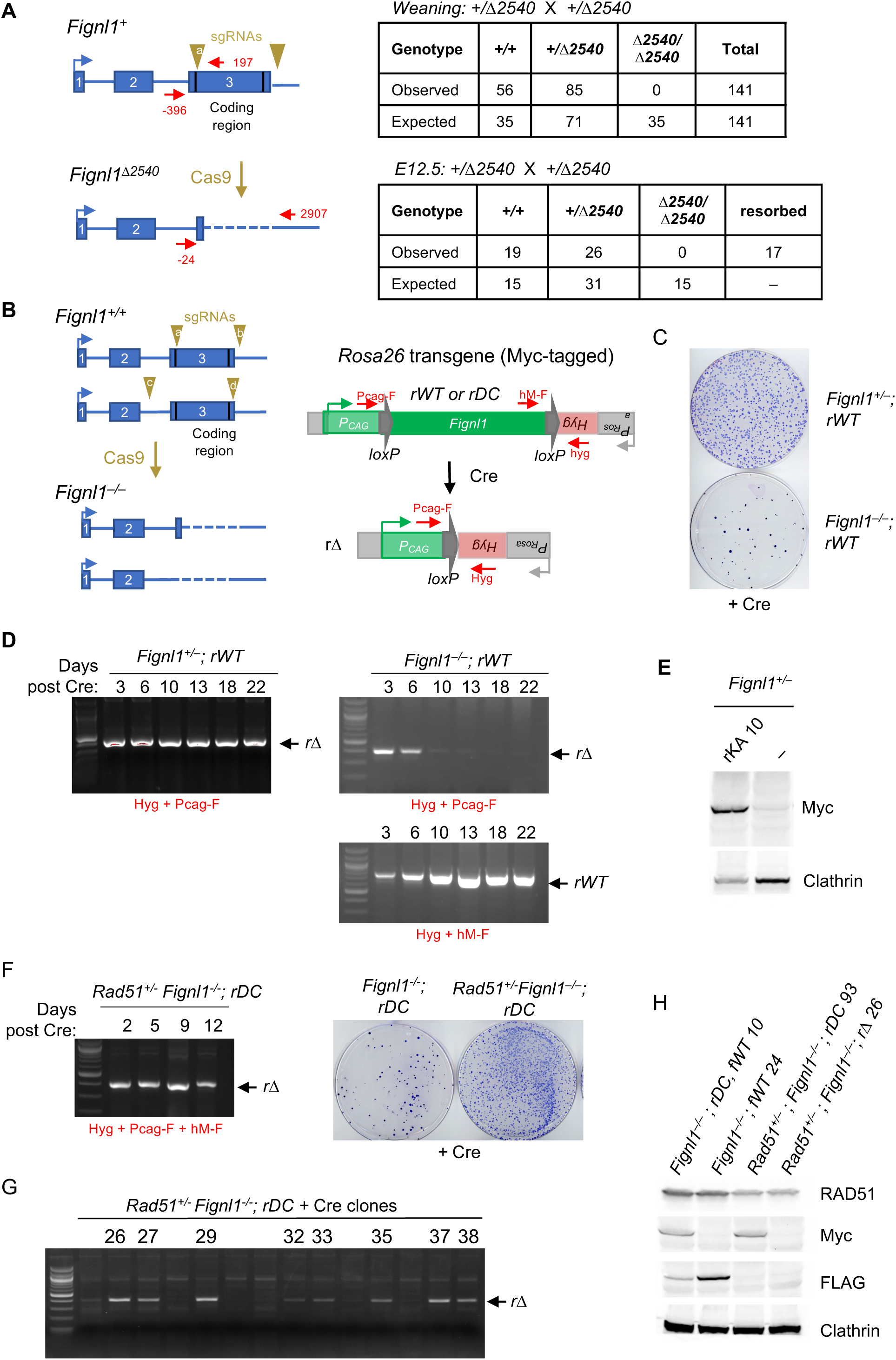
*Fignl1*^-/-^ mice and cells are not viable, but *Rad51* heterozygosity rescues cellular viability. (**A)** *Fignl1* mutant mice die prior to embryonic day 12.5. The *Fignl1^Δ2540^* allele was generated using CRISPR-Cas9 with two sgRNAs that delete the entire coding region. The position of the primers used for genotyping is shown relative to the ATG. No *Fignl1^Δ2540/Δ2540^* mutant mice were obtained either at weaning or at E12.5. (**B)** *Fignl1* conditional cell system. Wild-type *Fignl1* expressed from a *P_CAG_* promoter and flanked by loxP sites was targeted to the *Rosa26* locus (*rWT*). Correctly targeted cells were selected using *Hyg* expression from the *Rosa26* promoter, generating *Fignl1^+/+^*; *rWT* cells. The endogenous *Fignl1* alleles in the *Fignl1^+/+^*; *rWT* cells are deleted in two steps, first with sgRNAs (a+b) to delete one allele, generating *Fignl1^+/-^; rWT* cells, followed by sgRNAs (c+d) to delete the other allele, generating *Fignl1^-/-^; rWT* cells. Cre expression was then used to delete the wild-type *Fignl1* transgene *rWT* to generate the *rΔ* allele. Primers used to amplify either *rWT* or *rΔ* are indicated. **(C)** Reduced colony formation of *Fignl1^-/-^* cells in comparison to *Fignl1^+/-^* cells upon deletion of *rWT* upon Cre expression. **(D)** Similarly, loss of *rWT* results in impaired proliferation of *Fignl1^-/-^* cells. Genomic DNA was collected at the indicated days post transfection of the Cre expression vector and amplified by primers shown in **B**. While deletion at the *Rosa26* locus (*rΔ)* was readily detected at days 3 and 6 post-Cre expression in *Fignl1^-/-^; rWT* cells, it was not observed at later time points, indicating that *Fignl1^-/-^*; *rΔ* cells were out competed over time by cells that had not deleted *rWT*. **(E)** Western blotting showing Myc-tagged FIGNL1 K456A (KA) expression in *Fignl1^+^*^/−^*; rKA* cells used in Fig. 1A**. (F)** *Rad51* heterozygosity rescues cellular viability of *Fignl1* null mice. A *Fignl1* conditional allele was set up in *Rad51* heterozygous cells containing the *DC* allele at the *Rosa26* locus (*rDC*). Deletion of *rDC* was observed at late time points after Cre expression (9 and 12 days), indicating that *Fignl1^-/-^* cells are viable with *Rad51* heterozygosity. The high frequency of colony formation post Cre expression supports that *Rad51 ^+/-^ Fignl1^-/-^; rΔ* cells are viable. **(G)** Individual *Rad51^+/-^ Fignl1^-/-^* ; *rΔ* colonies were identified from those picked from the plate shown in **F**. **(H)** Reduced RAD51 and absent Myc-tagged FIGNL1 in a clone of *Rad51^+/-^ Fignl1^-/-^ ; rΔ* cells.

**Fig. S2.**
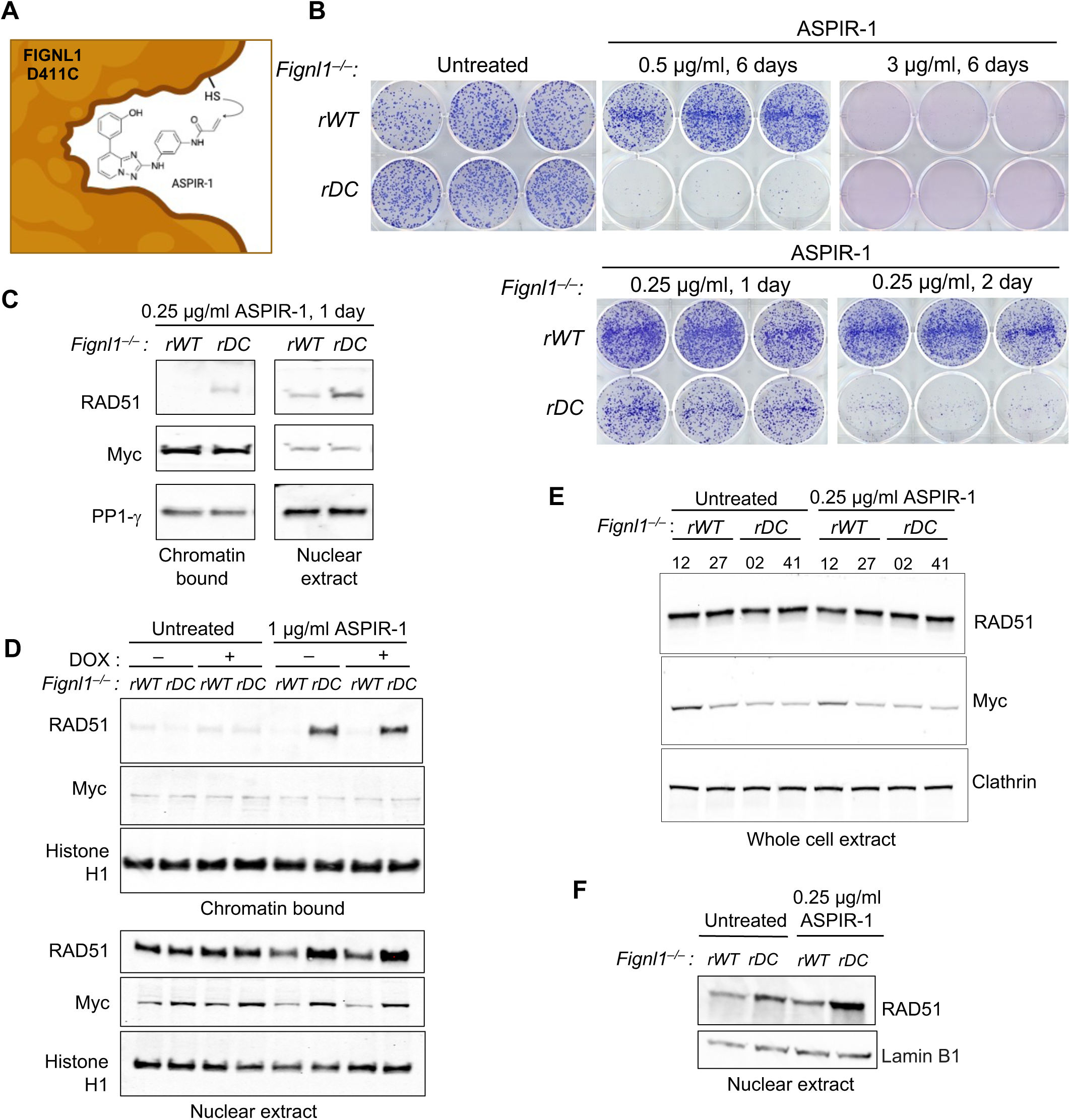
Chemical inhibition of FIGNL1 ATPase domain leads to cell lethality and is associated with RAD51 chromatin accumulation. **(A)** Chemical inhibition of mouse FIGNL1 D411C with ASPIR-1. ASPIR-1 is N-(3-((8-(3-hydroxyphenyl)-[1,2,4]triazolo[1,5-a]pyridin-2-yl)amino)phenyl)acrylamide (Cupido et al). The sulfhydryl group of the introduced in cysteine in the FIGNL1 catalytic site forms a covalent linkage with the acrylamide electrophile in ASPIR-1. ATPase activity of the cognate human FIGNL1 mutant D402C is reduced in a dose-dependent manner by ASPIR-1; in the absence of ASPIR-1, the D402C mutant has a 2-fold reduction in catalytic activity. The drawing was created with BioRender software. (**B)** Titration of ASPIR-1 in *Fignl1^-/-^; rWT* and *Fignl1^-/-^; rDC* cells. Continuous ASPIR-1 exposure at 0.5 µg/ml leads to lethality only of *Fignl1^-/-^; rDC* cells, whereas 3 µg/ml is also lethal to control cells. One day exposure to 0.25 µg/ml ASPIR-1 is sub-lethal, whereas 2-day exposure kills most *Fignl1^-/-^; rDC* cells. (**C)** A sublethal dose of ASPIR-1 is sufficient to lead to RAD51 accumulation in the chromatin fraction of *Fignl1^-/-^; rDC* cells. Here and below the *rWT* and *rDC* alleles encode Myc-tagged FIGNL1. **(D)** Similar to **C** with additional conditions. Cells were precultured for 3 days with doxycycline (1 µg/ml) where indicated to suppress BLM expression (Yusa et al). Western blot analysis of chromatin or nuclear fractions demonstrates similar level of RAD51 chromatin accumulation with or without BLM. **(E)** Total RAD51 levels are unaffected by ASPIR-1 treatment. Whole cell extracts from two independent clones of indicted genotypes were subjected to western blot analysis with the indicated antibodies. **(F)** Nuclear extract from cellular fractionation from Fig. 1C is shown. Three independent clones 02, 10, 41 of *Fignl1^-/-^; rDC* accumulated chromatin bound RAD51 in **C**, **D**, and **F,** respectively, in the presence of ASPIR-1.

**Fig. S3.**
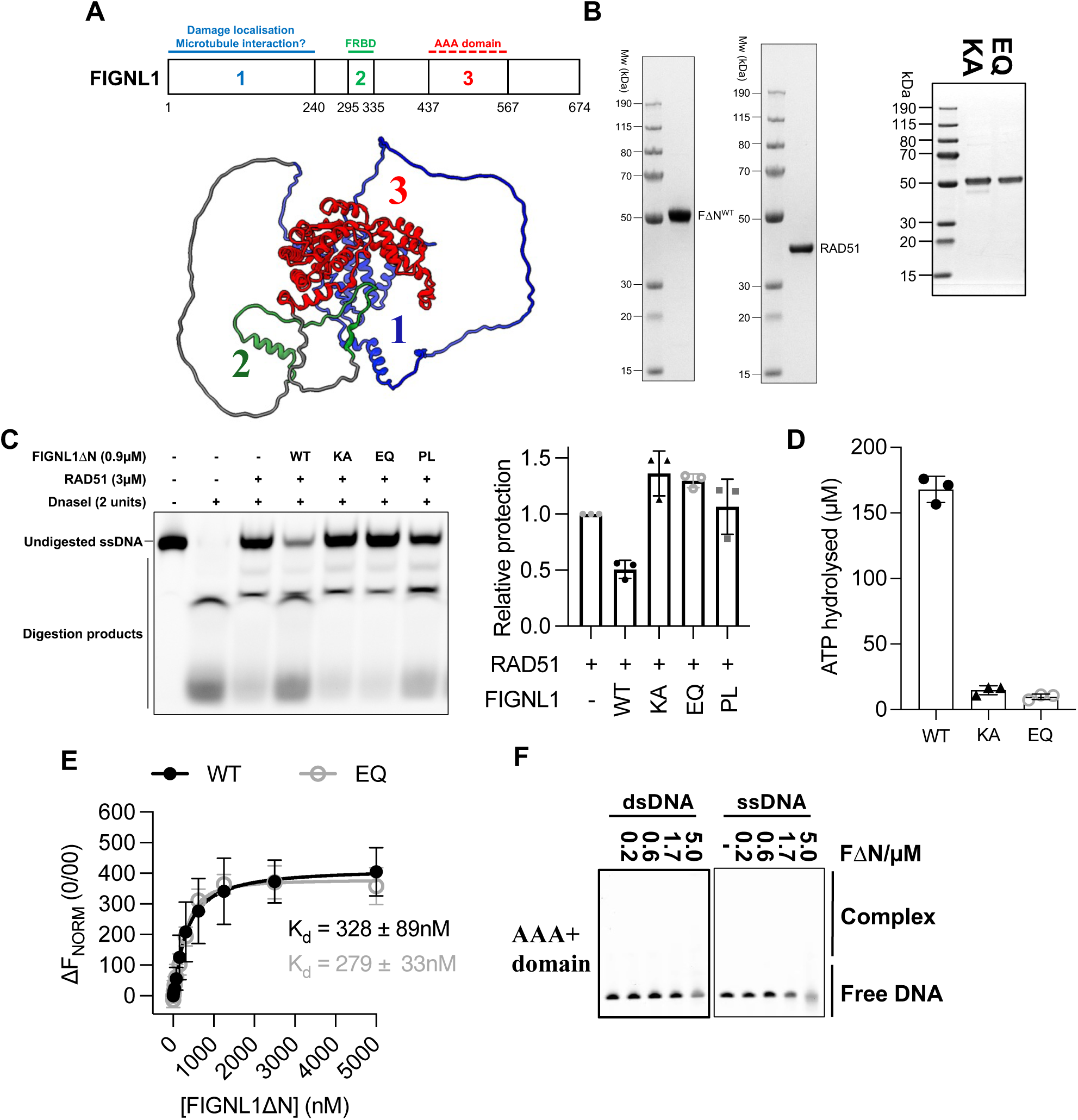
In vitro characterization of FIGNL1ΔN and mutants. **(A)** Domain architecture of FIGNL1 and AlphaFold2 model of full-length FIGNL1. **(B)** Images of gel showing quality of purified FIGNL1ΔN, RAD51, K447A and E501Q FIGNL1ΔN. **(C)** Image of gel showing nuclease protection of 60nt ssDNA by RAD51 in the presence of 0.9µM WT, Walker A – K447A, Walker B – E501Q and PL mutant of FIGNL1ΔN, and quantification of those nuclease protection assays. Each data point represents the mean ± s.d., n = 3. **(D)** Quantification of ATPase activity of E501Q, K447A and WT FIGNL1ΔN, n = 3, each data point represents the mean ± s.d. **(E)** Quantification of the interactions between WT and E501Q FIGNL1ΔN with RAD51 by MST, n = 3, each data point represents the mean ± s.d **(F)** EMSA data showing that the AAA+ domain of FIGNL1 does not bind to dsDNA or ssDNA.

**Fig. S4.**
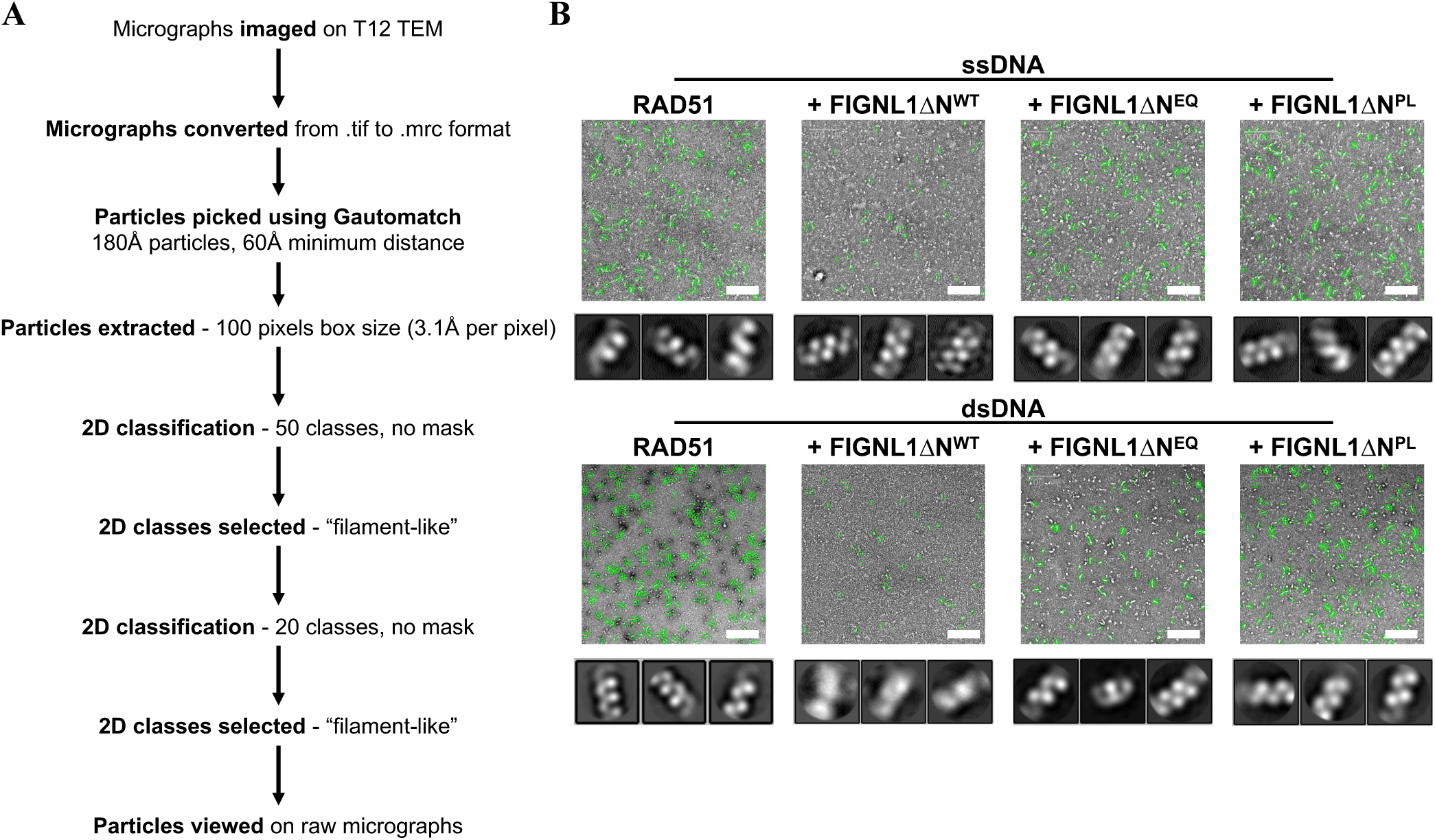
RAD51 filament quantification. (**A**) Processing pipeline of RAD51 filament length in the absence and presence of WT, E501Q and PL mutant FIGNL1ΔN. Briefly, micrographs were collected on a T12 TEM and subjected to particle picking using Gautomatch. Particles were extracted and subjected to two rounds of 2D classification after which the best “filament-like” particles were selected. The particles were checked visually and analysed using GraphPad Prism. (**B**) Representative micrographs and 2D class averages from each condition imaged. Scale bar = 200nm.

**Fig. S5.**
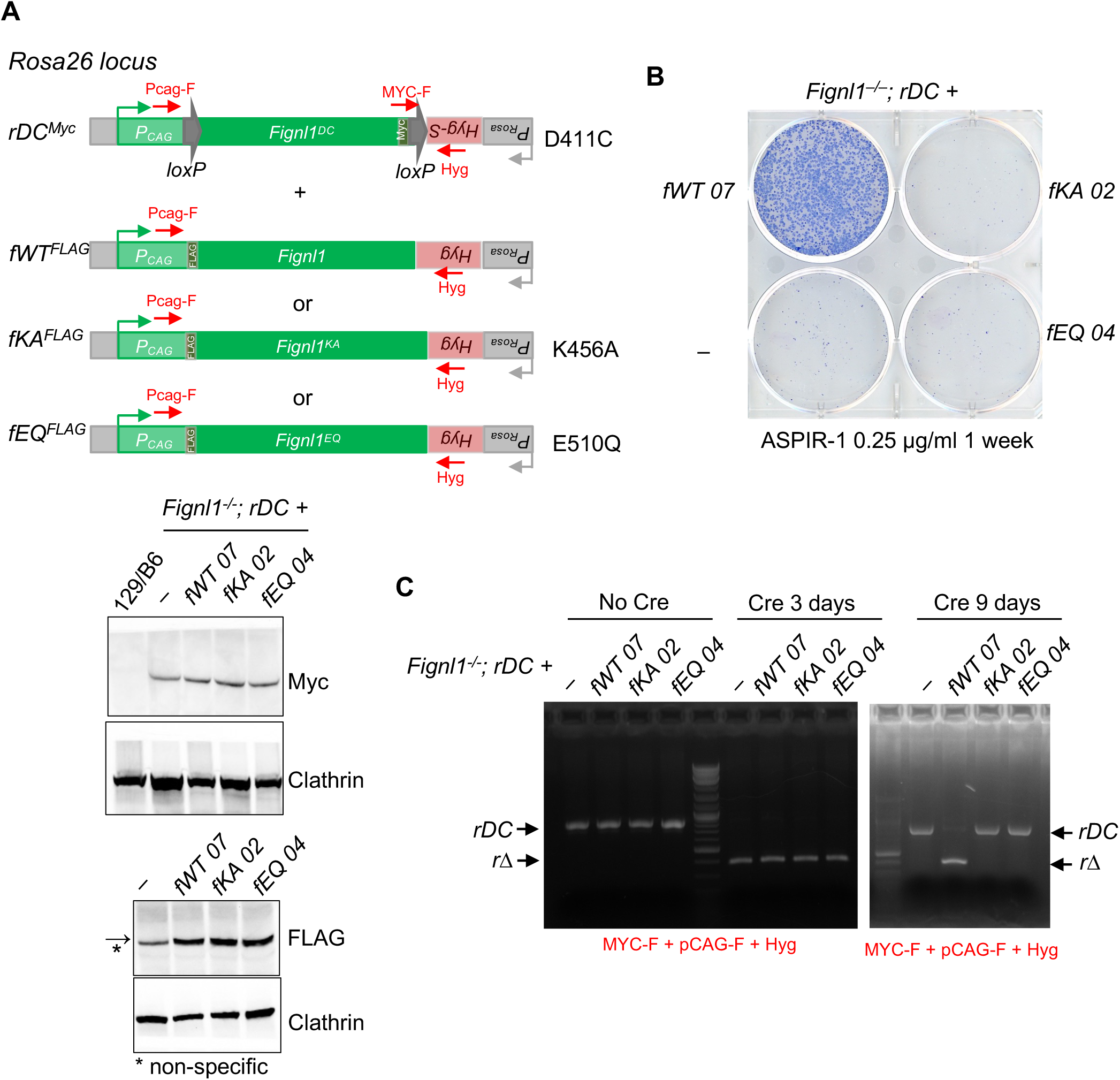
FIGNL1 Walker A and Walker B mutants are not able to suppress cellular lethality upon loss or chemical inhibition of the *rDC* allele, as seen using a dual *Rosa26* expression system. (A) The dual system is set up in three steps. 1. Integration of the conditional Myc-tagged *rDC* allele in one *Rosa26* allele, 2. disruption of the *Hyg* marker (*Hyg-S*, or sensitive), 3. introduction of a series for FLAG-tagged alleles at the other *Rosa26* allele. Western blotting confirms expression of the alleles. (B) *Fignl1^−/−^; rDC^Myc^, fWT^FLAG^* clones were resistant to ASPIR-1, indicating that the inhibited DC protein does not interfere with functional FIGNL1 protein. However, *Fignl1^−/−^; rDC^Myc^, fKA^FLAG^* or *Fignl1^−/−^; rDC^Myc^, fEQ^FLAG^* clones do not survive ASPIR-1 treatment, indicating that the Walker A and Walker B mutants, respectively, are unable (non-functional) to promote survival when the DC protein is inhibited. (**C)** Deletion of the *rDC* allele (*rΔ*) is stable in cells expressing wild-type FIGNL1 but not in cells expressing the Walker A and Walker B mutants, indicating that the *fKA* or *fEQ* expression impairs proliferation of cells. Genomic DNA was collected at the indicated days post transfection of the Cre expression vector and amplified by primers as shown in **A**.

**Fig. S6.**
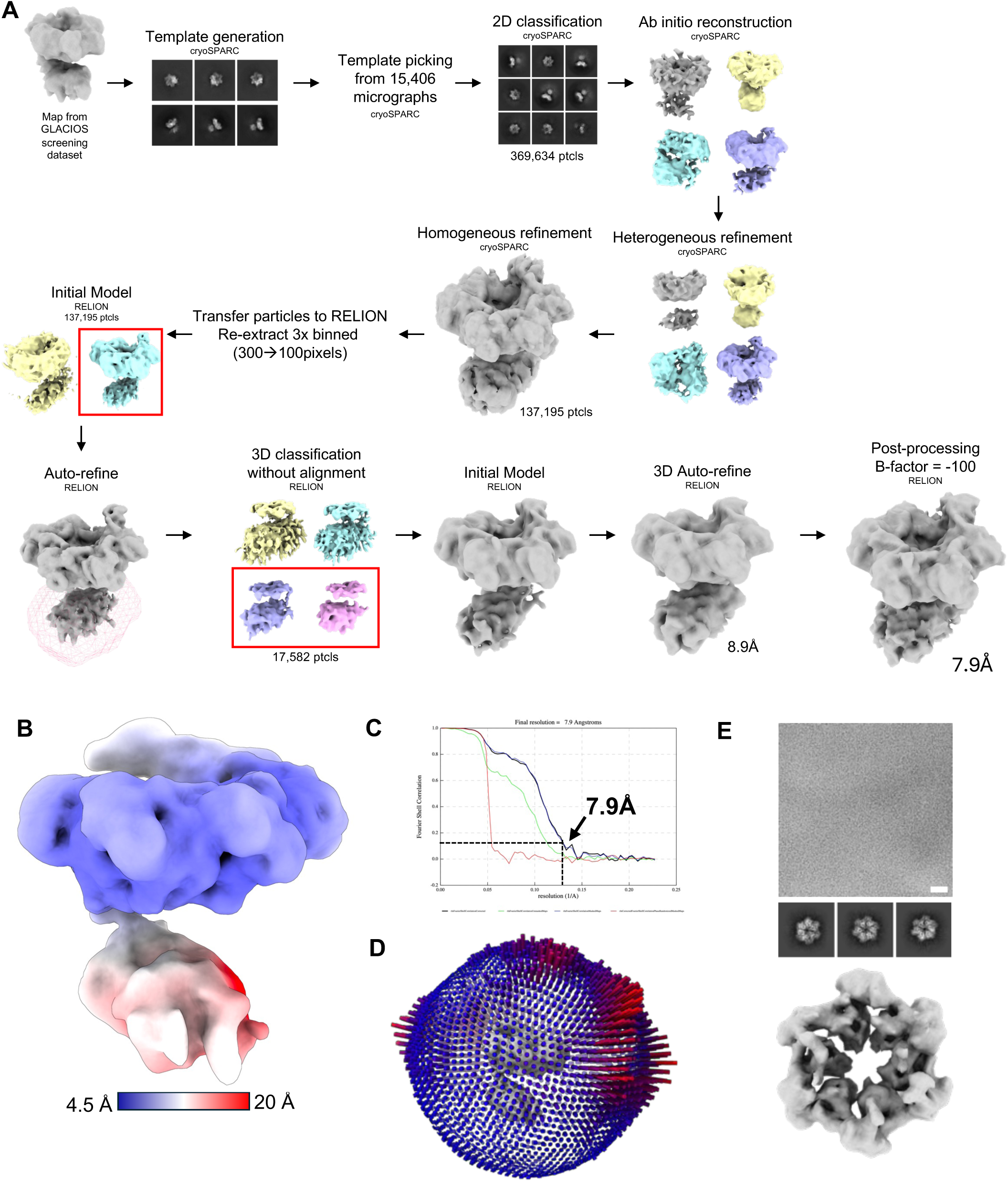
Single-particle data analysis flowchart for the FIGNL1ΔN^E501Q^-RAD51-ATP.Mg^2+^ complex. (**A**) Summary of the data processing pathway including the 2D classes, 3D model generation, 3D model selection, 3D classification and 3D refinement, resulting in the final reconstruction at 7.7 Å global resolution. (**B**) Local resolution distributions for the FIGNL1ΔN^E501Q^-RAD51 cryoEM reconstruction. **(C)** Fourier shell correlation (FSC) plots. **(D)** Angular distribution of particles contributed to the final reconstruction. **(E)** Micrograph, 2D classes and 3D reconstruction of WT FIGNL1ΔN in the presence of ATPγS, which shows that it can form hexamers. FIGNL1ΔN was applied to lacey carbon copper grids with an ultrathin carbon layer and imaged using on a KRIOS TEM. Particles were picked, 2D classified, refined and post-processed to produce a map of ∼10Å resolution. Scale bar = 50nm.

**Fig. S7.**
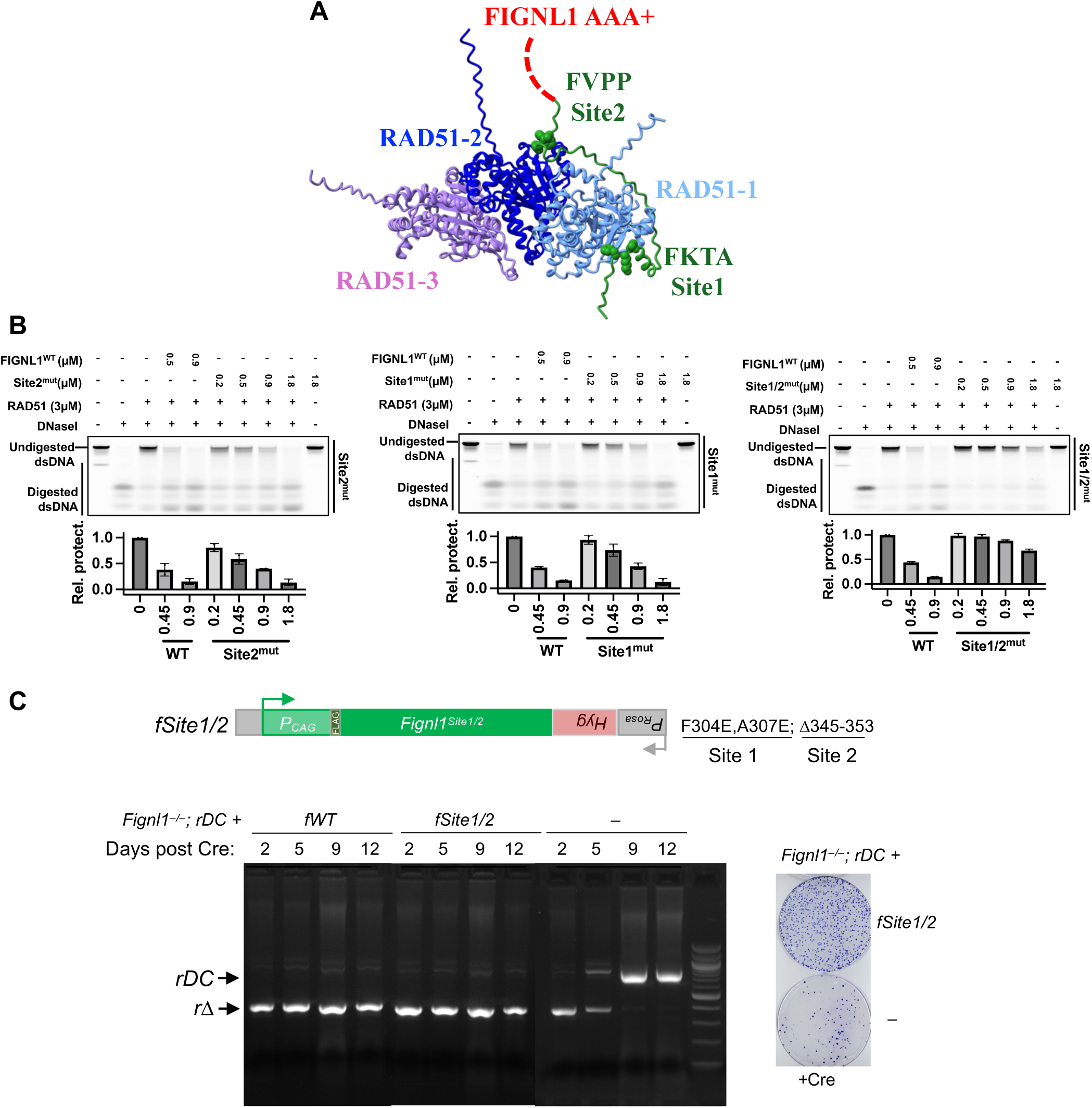
Probing the interactions between FIGNL1 and RAD51. **(A)** AlphaFold2 model of FIGNL1 FRBD in complex with a RAD51 trimer. **(B)** Nuclease protection assay showing the effect of WT FIGNL1ΔN, Site1 deleted (Site1^mut^), Site2 deleted (Site2^mut^) and both sites deleted (Site1/2^mut^) formed on 60mer dsDNA coated by RAD51. Quantification was shown underneath. Each data point represents the mean ± s.d. n = 3. (**C**) The FLAG-tagged Site1/2 FIGNL1 mutant F304E, A307E, Δ345-353 mutant was expressed from the second allele in the dual *Rosa26* system (*Fignl1^−/−^; rDC, fSite1/2*). Deletion of the *rDC* allele is detected at days 9 and 12 post-Cre transfection in *Fignl1^−/−^; rDC fSite1/2* or *fWT* cells, while it is only initially observed in *Fignl1^−/−^; rDC* cells and then is gone by the days 9 and 12, indicating that the Site1/2 mutant is still functional for supporting cells viability. This is supported by colony formation shown in the right panel. *Fignl1^-/-^; rΔ* cells were unable to form colonies, whereas *Fignl1^−/−^; rΔ, fSite1/2^FLAG^* cells robustly formed colonies.

**Fig. S8.**
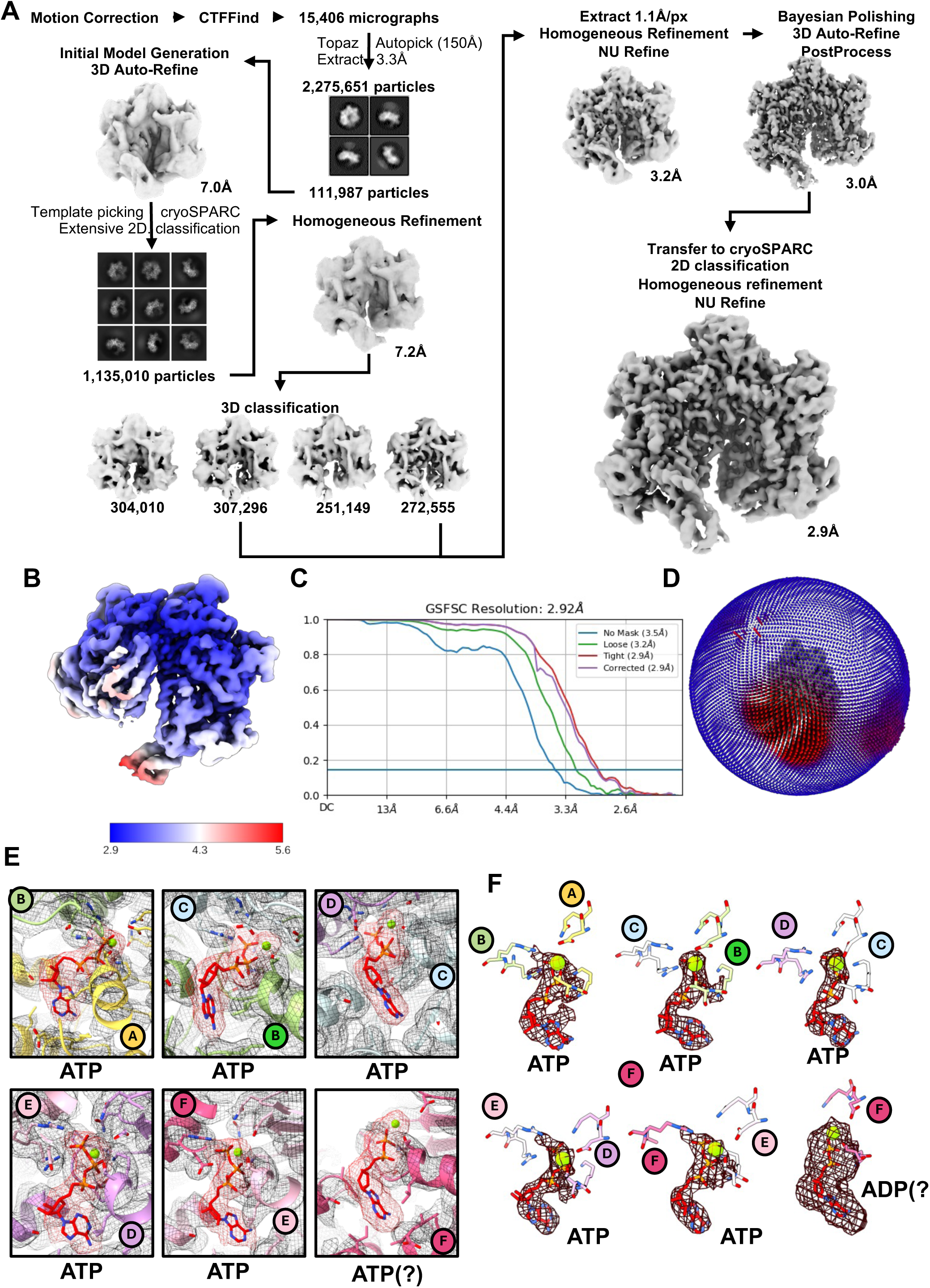
Single-particle analysis (SPA) of the FIGNL1ΔN^E501Q^-RAD51 complex. **(A)** Data processing pipeline showing the pathway from raw movies to the final 2.9Å reconstruction of the FIGNL1ΔN hexamer bound to RAD51, focusing on FIGNL1 AAA+ domain. Classes taken to subsequent processing steps are outlined in red. **(B)** Local resolution estimates for the FIGNL1ΔN^E501Q^-RAD51 cryoEM map. **(C)** Fourier shell correlation (FSC) plots. **(D)** Angular distribution plot. (**E**) electron density and models surrounding each nucleotide binding pocket. (**F**) Density for nucleotide in each nucleotide binding pocket.

**Fig. S9.**
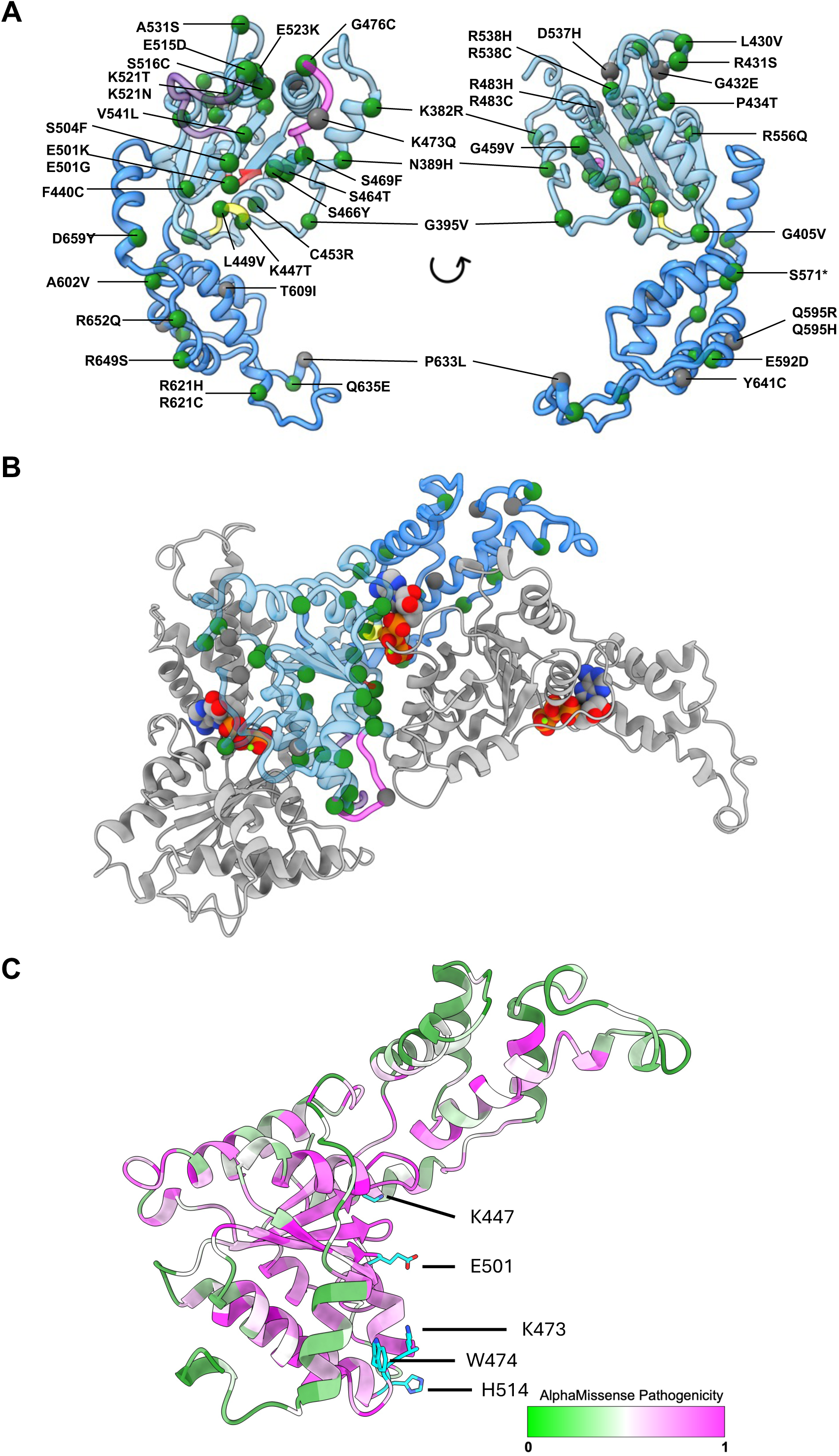
Mutations found in patients with cancer or various of unknown significance (VUS) in other genetic disorders in the FIGNL1 AAA+ domain. VUS are shown in gray, cancer mutations in green, pore loop 1 in magenta, pore loop 2 in purple, Walker A motif in yellow and Walker B in red. (**A**) Specific mutations mapped onto monomers (**B**) mutations in the context of oligomers. (**C**) Pathogenicity scores were calculated with AlphaMissense(39). Scores from 0 (low pathogenicity) to 1 (high pathogenicity) are colored on the AlphaFold model of the FIGNL1 AAA+ domain.

**Fig. S10.**
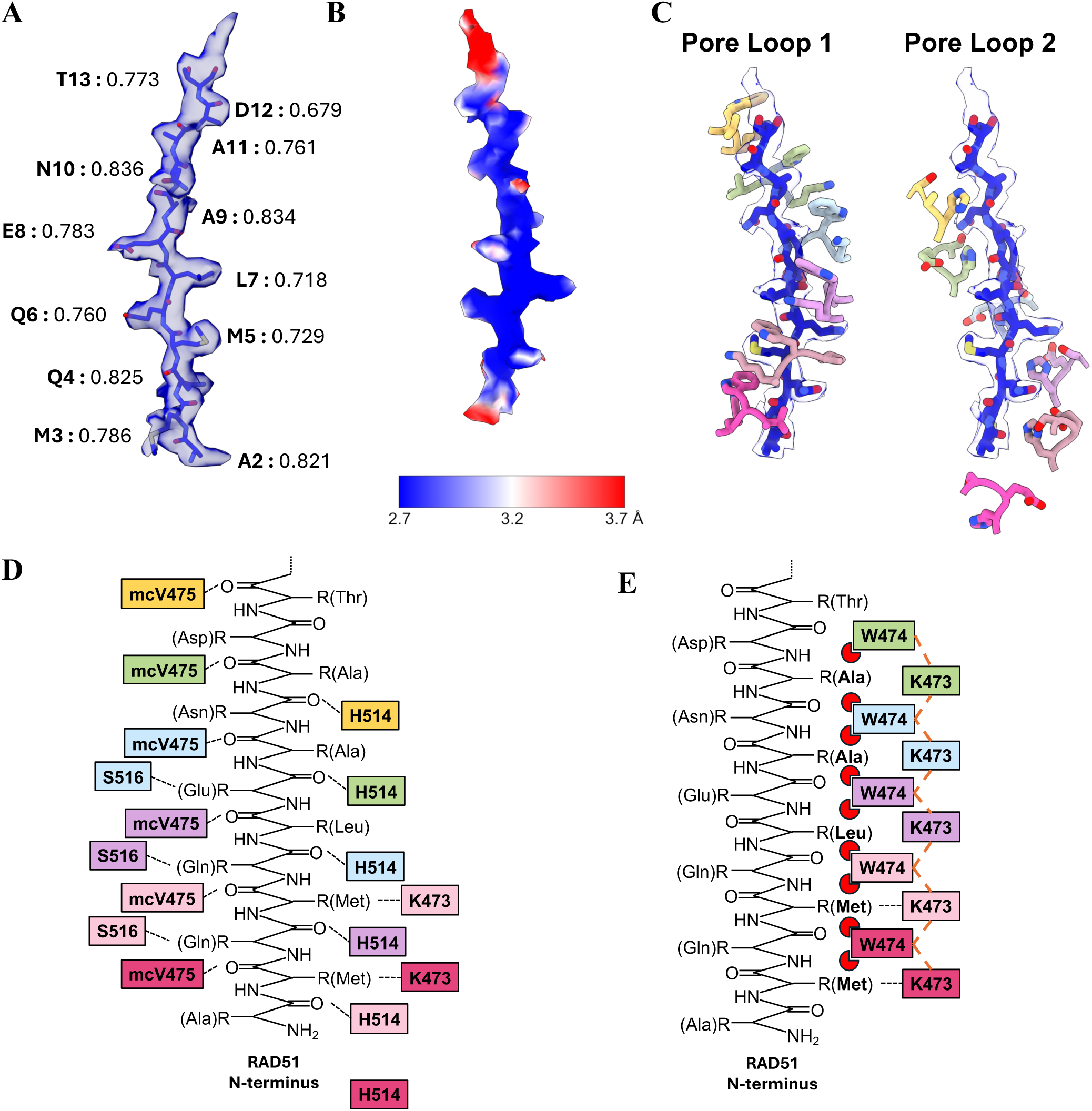
The pore loops of FIGNL1 form a close network of interactions with the RAD51 N-terminus. **(A)** Calculated Q-scores for the N-terminus of RAD51 fitted into the peptide density from the 2.9Å reconstruction of the peptide-bound FIGNL1 AAA hexamer [22]. **(B)** Local resolution estimates of the N-terminus of RAD51 as calculated in ResMap indicating high resolution of the N-terminus **(C)** The lysine and tryptophan residues of pore loop 1 of FIGNL1 intercalate the N-terminal peptide of RAD51. H514 of pore loop 2 potentially form hydrogen binding interactions with RAD51 N-terminal residues. **(D)** Schematic of putative H-bonds between pore loops 1 and 2 with the RAD51 N-terminus, main chain interactions are preceded by ‘mc’. **(E)** Schematic of the interaction network created by the lysine-tryptophan dipeptide of pore loop 1 and H514 of pore loop 2. Orange dashed lines represent π-cation interactions. Red circles represent hydrophobic interactions from W474 to the hydrophobic side chains of RAD51.

**Fig. S11.**
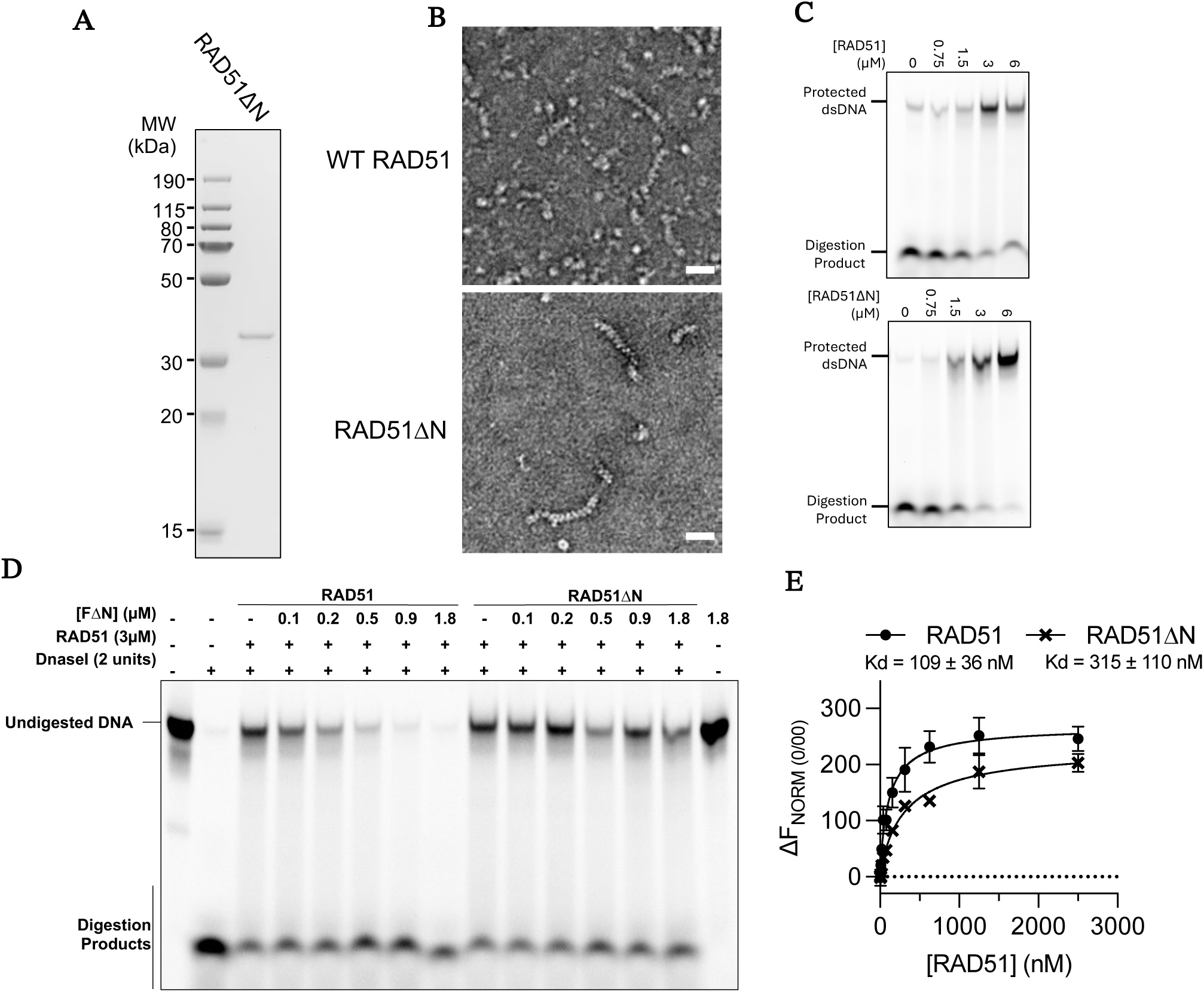
Characterisation of RAD51ΔN. **(A)** Image of gel showing quality of purified RAD51ΔN. **(B)** Representative micrographs of RAD51 and RAD51ΔN in complex with dsDNA. **(C)** Image of gel of nuclease protection of dsDNA by increasing concentrations of RAD51 and RAD51ΔN. **(D)** Image of gel of nuclease protection by RAD51 and RAD51ΔN in the presence of increasing concentrations of WT FIGNL1ΔN. **(E)** Quantification of interaction between labelled WT FIGNL1ΔN and WT RAD51 and RAD51ΔN as measured by microscale thermophoresis of AlexaFluor-647-labelled WT FIGNL1ΔN. Each datapoint represents the mean ± S.E.M, n = 3.

**Fig. S12.**
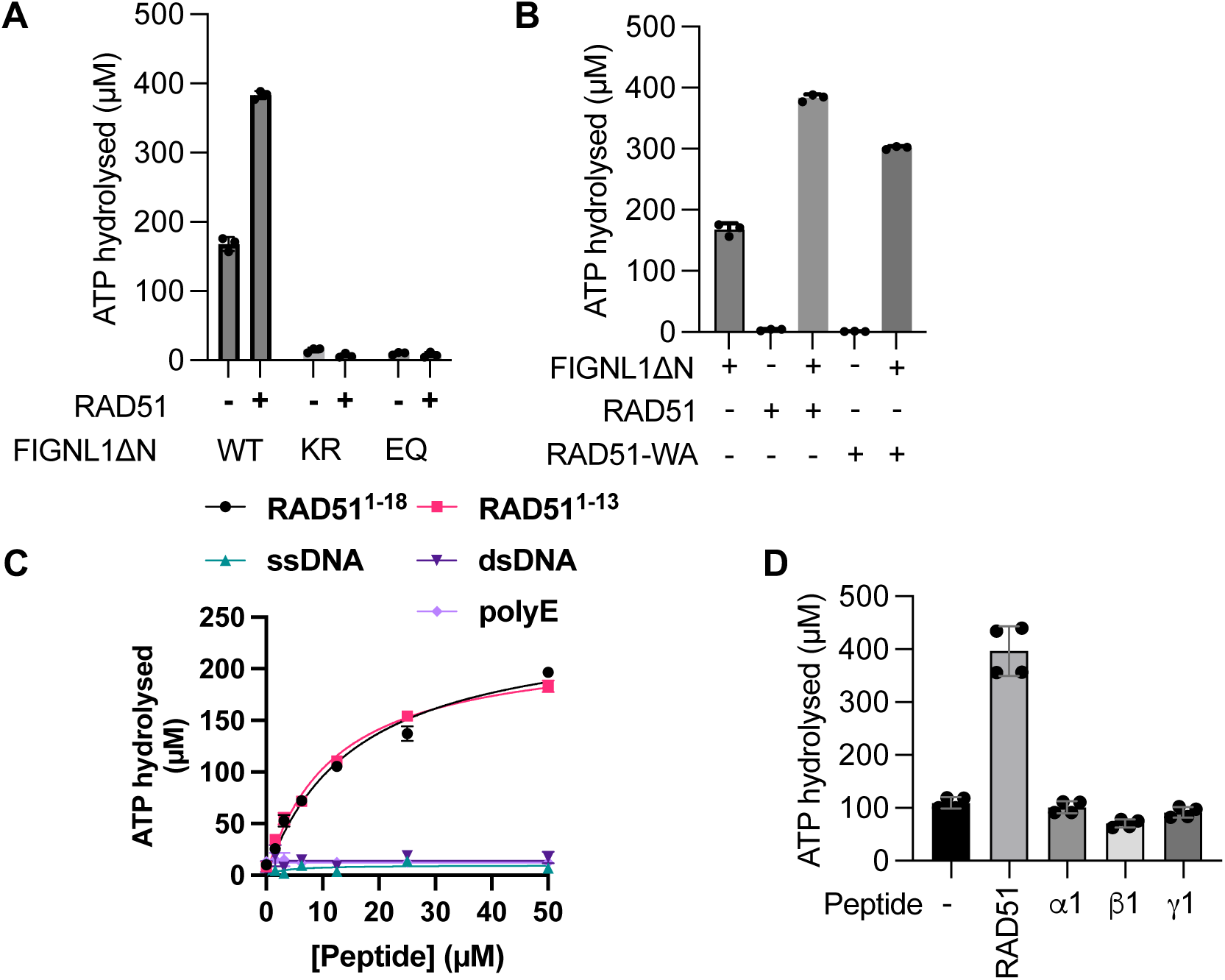
Investigation of the effect of RAD51 on FIGNL1 ATPase activity. (**A**) ATP hydrolysis rate of FIGNL1ΔN, Walker A (K477R), Walker B (E501Q) mutant in the absence and presence of RAD51. **(B)** ATP hydrolysis rate of FIGNL1ΔN in the absence and presence of WT RAD51 and RAD51 Walker A mutant (RAD51-WA, K133R). (**A**) and (**B**) together support that RAD51 stimulates FIGNL1 activities while FIGNL1 does not stimulate RAD51 ATPase activity. (**C**) ATP hydrolysis by FIGNL1ΔN in the presence of mutated RAD51 N-terminal peptide. **(c)** ATP hydrolysis by FIGNL1ΔN in the presence of RAD51 N-terminal peptides (1-18 and 1-13), 60mer ssDNA and dsDNA and polyglutamate peptide. Each data point represents 3 independent measurements. (**D**) ATP hydrolysis of FIGNL1ΔN (1 μM) in the presence and absence of 50µM RAD51 N-terminal peptide and α, β and γ tubulin C-terminal peptides. Each data point represents the mean ± standard deviation.

**Fig. S13.**
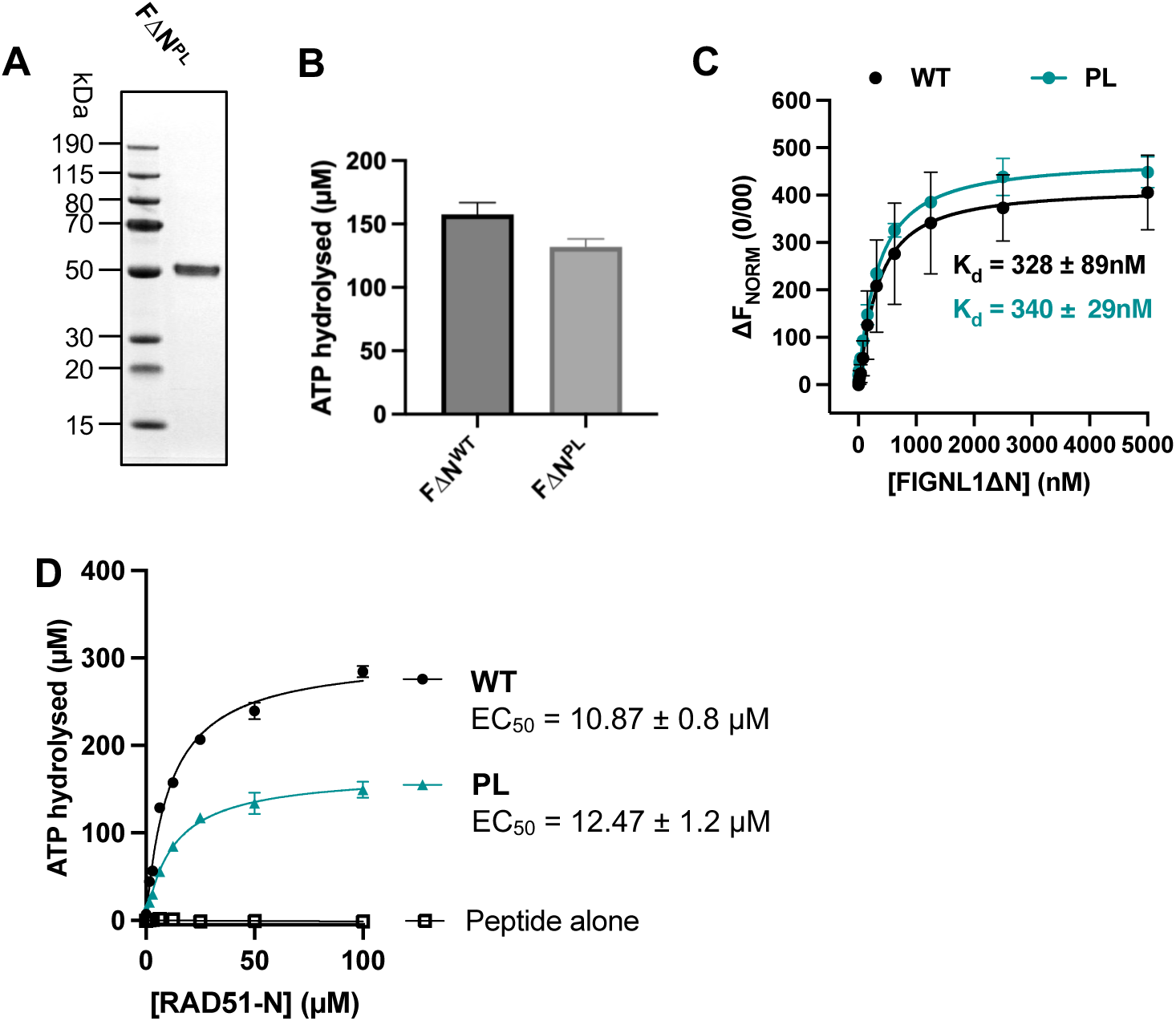
Characterization of the PL mutant of FIGNL1ΔN and investigating the translocase activities of FIGNL1. **(A)** Image of gel showing quality of purified PL mutant FIGNL1ΔN (K473E, W474A). **(B)** Quantification of the ATPase activities of WT and PL FIGNL1ΔN, n = 3, data points represent the mean ± s.d. **(C)** Quantification of the WT and PL FIGNL1ΔN interaction with RAD51 as measured by microscale thermophoresis, n = 3, data points represent the mean ± s.d. WT data shown for comparison but it is the same data as shown in Extended Data Figure 3e. (**D**) Quantification of the ATPase activity of 100nM WT and PL FIGNL1ΔN in the presence of increasing concentrations of RAD51 N-terminal (RAD51-N) peptide. Each data point represents the mean ± s.d. ATPase activity of the PL mutant can be stimulated by RAD51.

**Fig. S14.**
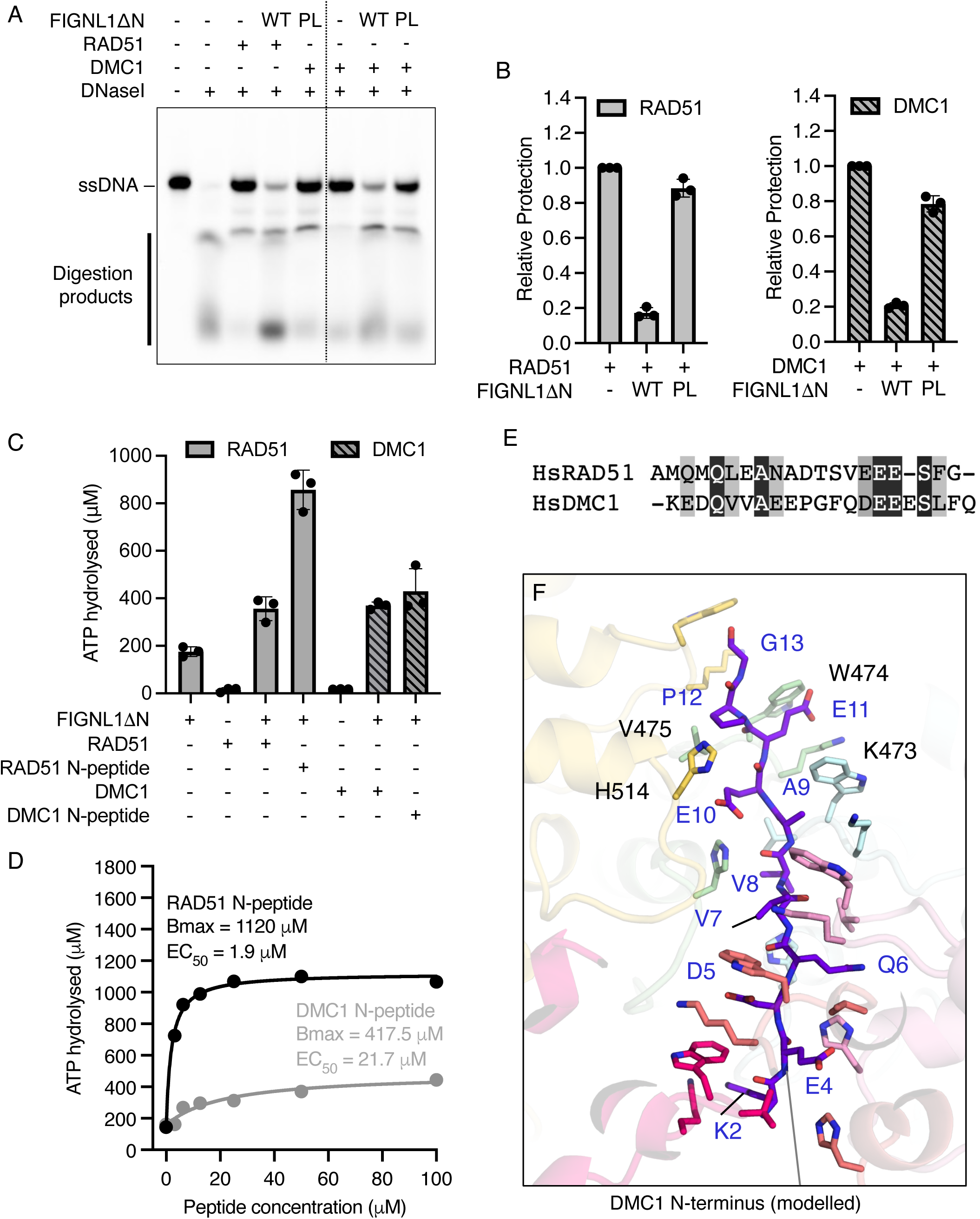
FIGNL1ΔN acts upon DMC1. **(A)** Image of polyacrylamide gel showing nuclease protection of 60nt ssDNA by RAD51 or DMC1 alone and in the presence of 0.9μM WT or PL (pore loop) FIGNL1ΔN. **(B)** Quantification of replicated nuclease protection experiments. Bar represents mean ± standard deviation. Individual data points for each replicate is shown. Relative protection is normalized to either RAD51 or DMC1 alone for the respective experiments. **(C)** ATP hydrolyzed by FIGNL1ΔN (1μM) in the absence and presence of 1μM RAD51, 50μM RAD51 N-terminal peptide, 1μM DMC1, or 50μM DMC1 N-terminal peptide. Bar represents mean ± standard deviation. Individual data points for each replicate is shown. **(D)** Stimulation of FIGNL1ΔN ATPase activity by RAD51 and DMC1 N-terminal peptides at 1μM FIGNL1. **(E)** Amino acid sequence alignment between the N-terminal regions of human RAD51 and DMC1. Shaded residues are similar (grey) or identical (black). **(F)** Modelling of the DMC1 N-terminus into the pore of FIGNL1, based upon the FIGNL1 hexamer-RAD51 cryoEM complex determined in this study, suggests it can be accommodated. Pore loop side chains are shown as sticks and subunits colored as for Fig 2.

**Fig. S15.**
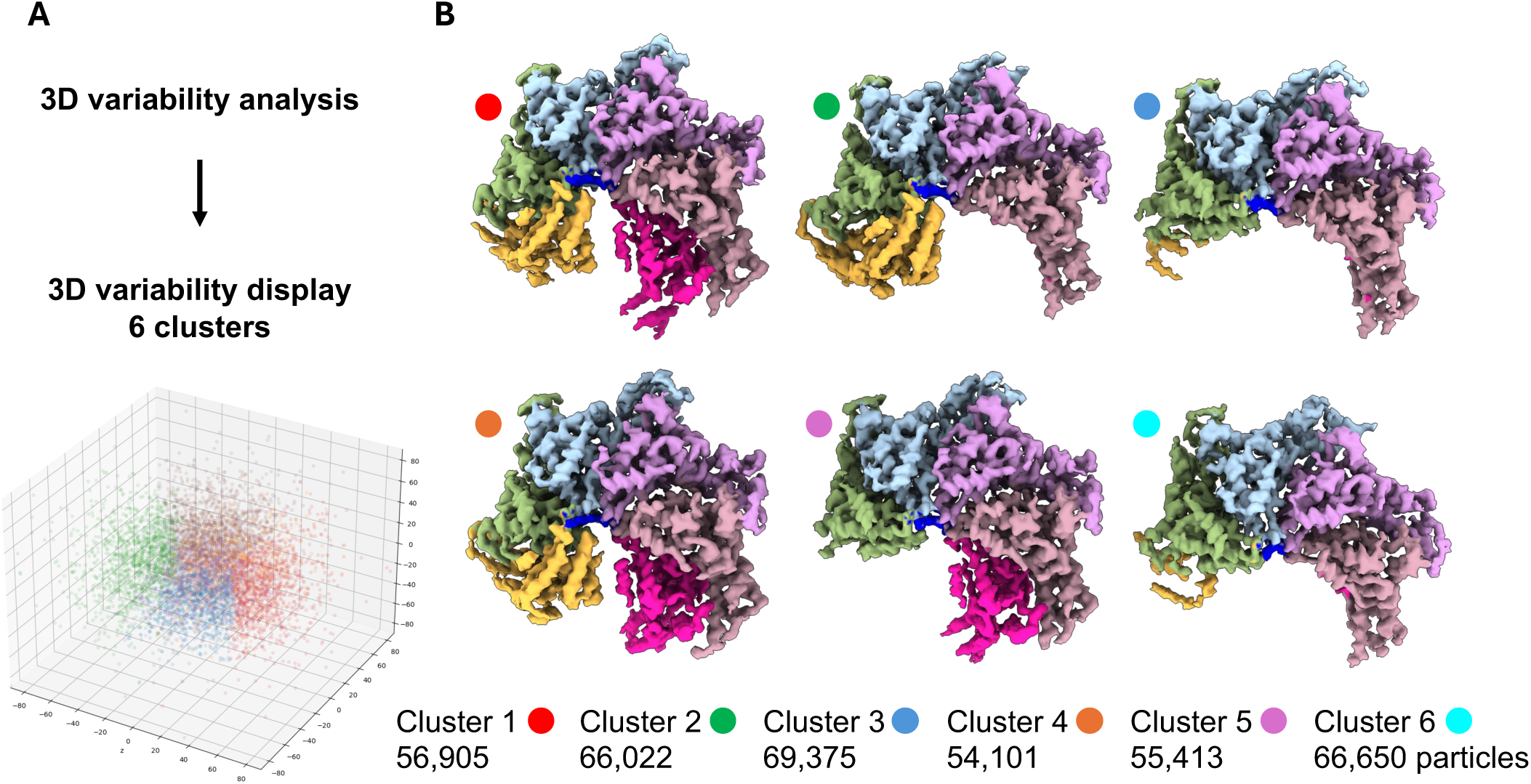
3D variability analysis of the FIGNL1 AAA+ hexamer bound to RAD51. **(A)** Processing pipeline showing source of particles that were used as an input for 3D variability analysis and graph of principle component analysis of the outputted clusters showing grouping of particles. **(B)** 3D volumes showing the tetrameric, pentameric and hexameric species present in the FIGNL1-RAD51 cryoEM dataset.

**Fig. S16.**
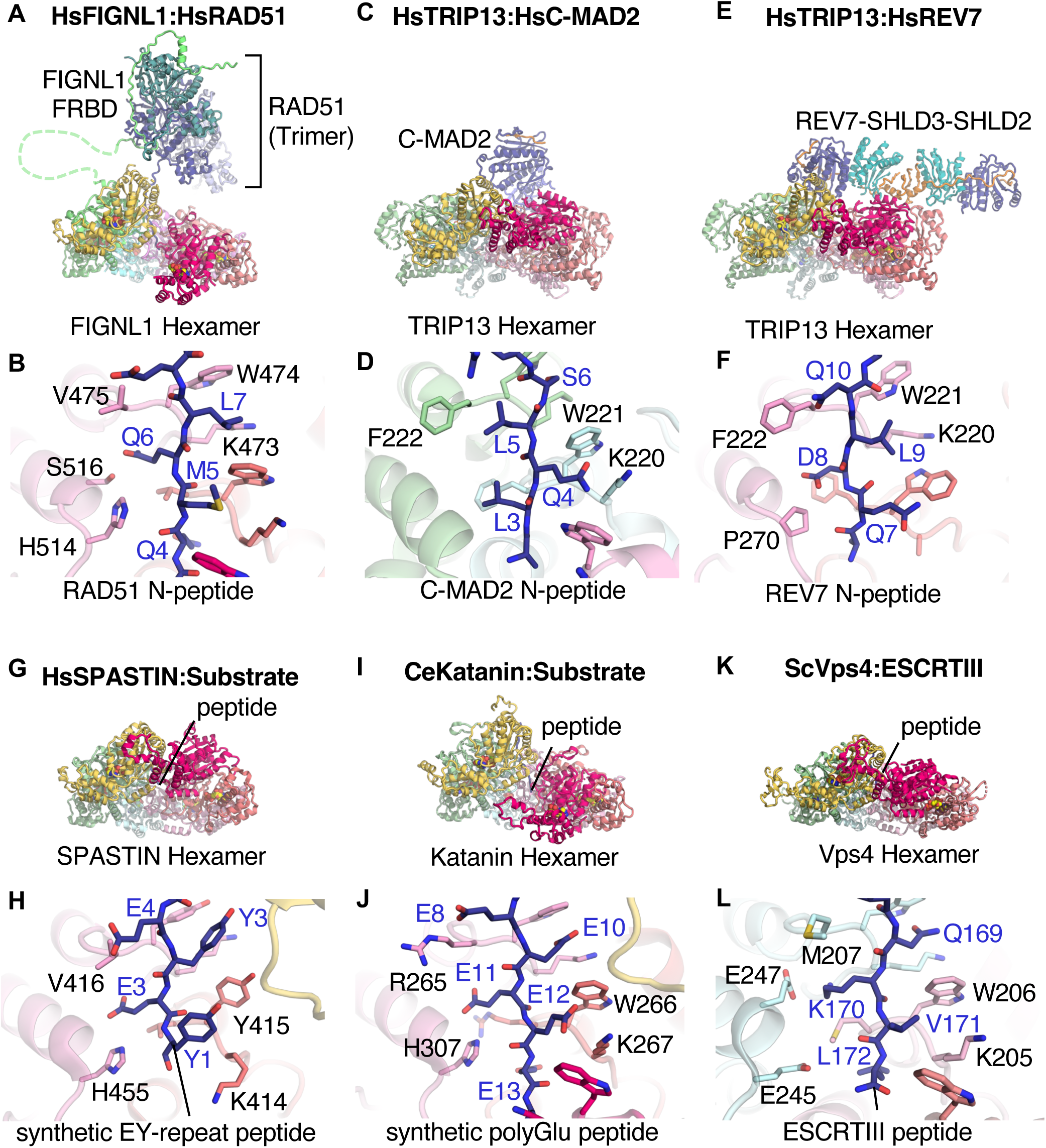
Structural comparisons between FIGNL1 and peptide translocation AAA ATPases. **(A)** Composite atomic model of human FIGNL1-RAD51 using the cryoEM structure in combination with the Alphafold2 multimer predictions of FIGNL1-FRBD in complex with a RAD51 trimer. **(B)** Molecular details of the KWV-containing pore loop 1, and contributions from pore loop 2, interacting with the RAD51 N-terminus. **(C)** Structure of human TRIP13 bound to C-MAD2 (PDB 6F0X) alongside **(D)** the molecular details of the KWF-containing pore loop 1 interacting with the N-terminus of C-MAD2. **(E)** Structure of human TRIP13 bound to REV7 complex (PDB 7L9P) with **(F)** the molecular details of the pore loops interacting with REV7 substrate. **(G)** Structure of human SPASTIN in complex with a synthetic peptide glutamate-tyrosine dipeptide repeat substrate (PDB 6PEN) with the molecular details of the KYV-containing pore loop 1 interacting with the substrate **(H)**. **(I)** Structure of *C. elegans* Katanin in complex with a synthetic glutamate-rich peptide (PDB 6UGD) with the molecular details of the KWR-containing pore loop interacting with the substrate **(J)**. **(K)** Structure of *S. cerevisiae* Vps4 in complex with an ESCRTIII-peptide substrate (PDB 6AP1) with the molecular details of the KWM-containing pore loop 1 interacting with the peptide **(L)**. Structures were rendered in PyMol (Schrodinger). All structures are aligned with respect to the FIGNL1 hexamer and the subunits coloured by protomer (see Fig 3.) Substrates found threaded through the AAA+ hexamer pore are coloured dark blue.

**Fig. S17.**
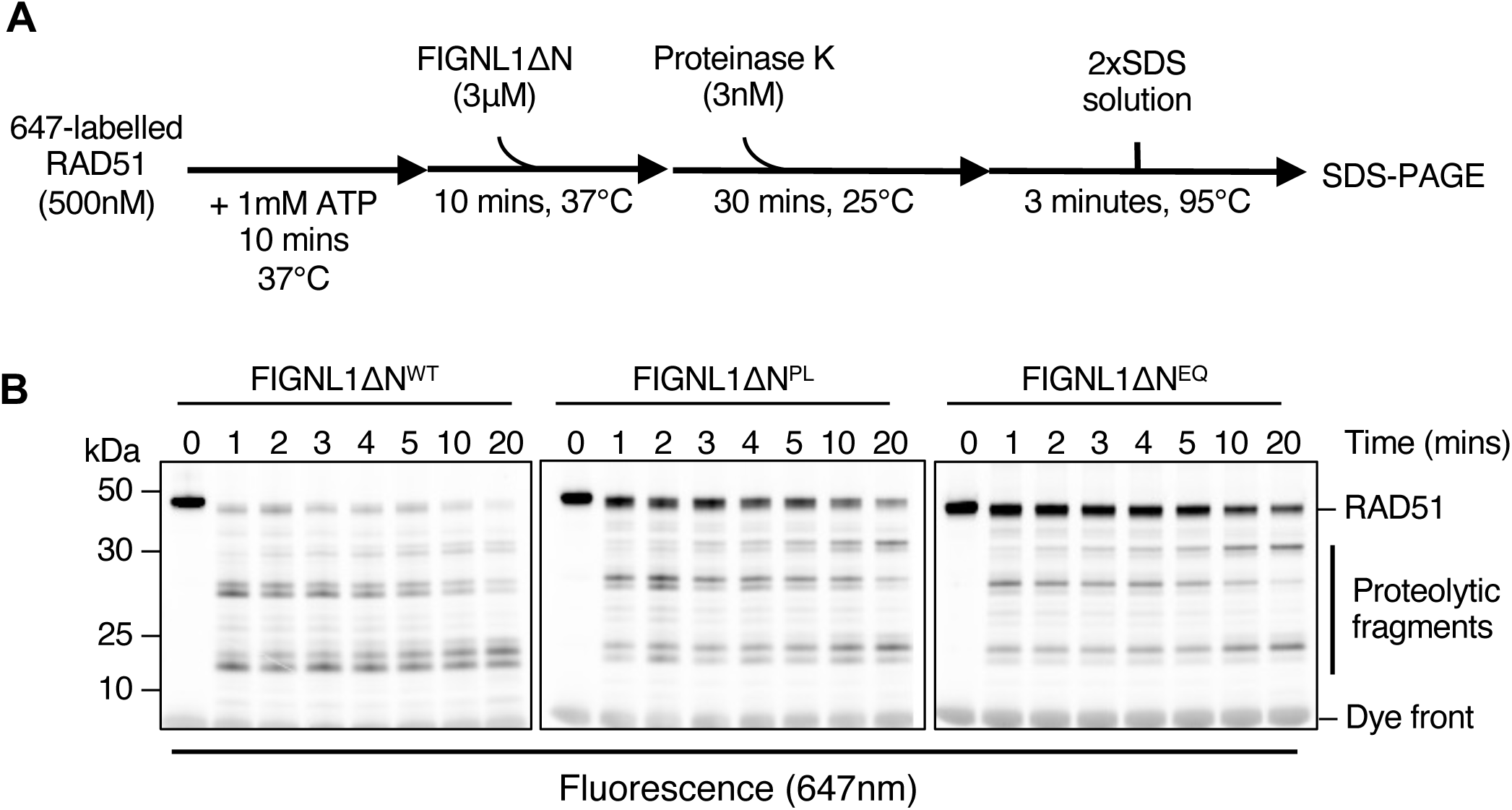
Probing RAD51 conformation using limited proteolysis. (**A**) Limited proteolysis experiment scheme. (**B**) Proteinase K digestion of RAD51 in the presence of a 6-fold molar excess of either wild-type (WT), pore-loop mutant (PL), or Walker B (EQ) mutant FIGNL1ΔN at 25°C. Proteins and proteolytic fragments were resolved by 4-12% SDS-PAGE. RAD51-specific protein bands were visualised using a covalently attached fluorescent label (647nm). Each lane represents samples withdrawn at indicated time points. DNA was omitted in these experiments to remove conflating proteolytic patterns from filament to monomer transitions.

